# Chromosome-scale comparative genomics reveals a genomic mosaic across the documented Hattan sake-brewing rice lineage

**DOI:** 10.64898/2026.11.23.765307

**Authors:** Ryo Nozu, Zennosuke Katsuba, Miyuki Kanda, Ayane Fujita, Sora Yonezawa, Shuji Matsushita, Hidemasa Bono

## Abstract

**Background:** Breeding pedigrees record crosses and selections but do not reveal how the documented relationships are reflected in descendant genomes. Therefore, whole-genome summaries can mask local inheritance patterns generated by recombination and segregation. To examine how documented relationships are reflected across genomic scales, we generated chromosome-scale assemblies of the Hattanso landrace and four cultivars from the documented Hattan sake-brewing rice lineage of Hiroshima, Japan. We then analyzed genome-wide and local relationships using small variants, structural variants, and gene presence–absence variation.

**Results:** The five assemblies ranged from 375.78 to 378.94 Mb in length, with scaffold N50 values of 30.25–30.43 Mb and 99.8% BUSCO completeness. Single-nucleotide variant data identified Hattan Nos. 10 and 35 as the closest pair at the genome-wide scale, a relationship also supported by protein sequence alignment. This relationship was recovered on 9 of the 12 chromosomes, but genomic relationships varied among local regions. Classification of 500-kb windows based on predefined allele-sharing patterns associated with documented breeding steps assigned 51.91% of the nuclear genome to lineage-pattern categories and identified 143 candidate transition intervals. Publicly available reads from Shuho broadly supported the Hattan No. 35 state, consistent with the recorded Hattan No. 10 × Shuho parentage of Hattan No. 35. Hattan-nishiki Nos. 1 and 2, selected from the recorded Hattan No. 35 × Akitsuho cross, exhibited distinct genomic configurations, with differences across all three variant classes concentrated on Chr12. A 1-Mb region on Chr4 contained single-nucleotide variants, small indels, and structural variants predominantly shared by Hattanso and the four derived cultivars. Functional enrichment analysis of lineage-associated gene sets identified pectinesterase-related functions in the Hattan No. 10-specific set and a diterpenoid-related signal in the Hattan-nishiki No. 2-specific set.

**Conclusions:** The documented Hattan sake-brewing rice lineage exhibits a spatially heterogeneous genomic mosaic, in which a strong genome-wide relationship coexists with local variation in lineage-associated states. Integrating nucleotide-scale, structural, and gene-content variation across genomic scales thus complements pedigree records and helps prioritize candidate regions and genes for future functional and breeding studies.

## Background

Crop breeding pedigrees record crosses and selections but do not reveal how ancestral genomic segments are distributed among descendant cultivars. Recombination and segregation reorganize inherited chromosomal segments through successive generations. Thus, even cultivars with straightforward documented breeding relationships can differ in the genomic material available for selection. In Japanese rice, genome-wide single-nucleotide polymorphism (SNP) analyses have resolved pedigree haplotypes and traced chromosomal block transmission through historical breeding (Yamamoto et al. 2010). Recently, an SNP-based haplotype visualization method has been used to reconstruct chromosomal block transmission and estimate recombination regions within a modern rice lineage (Kwon et al. 2025). Together, these studies demonstrate the value of integrating pedigree records with chromosome-level inheritance patterns.

The Hattan rice lineage in Hiroshima provides a well-documented system for investigating genomic inheritance across successive breeding stages. Hattanso is the local landrace from which the documented Hattan lineage originated, and Hattan No. 10 was developed from Hattanso through pure-line selection (Hiroshima Prefectural Agricultural Technology Center 1999, p. 22). Hattan No. 35 was subsequently developed by crossing Hattan No. 10 with Shuho, followed by selection (Takei et al. 1968). Hattan-nishiki Nos. 1 and 2 were later selected as sister cultivars from a cross between Hattan No. 35 and Akitsuho (Maeshige et al. 1984a, b). Previous studies have characterized the grain properties, polishing performance, and sake-brewing suitability of Hattan-lineage materials (Tamaki et al. 2005a; Tamaki et al. 2005b; Tamaki et al. 2007). Together, these breeding records and phenotypic studies provide a foundation for genomic comparisons across successive breeding stages.

Analyzing structural and gene-content variation alongside SNP-defined haplotype blocks can provide a more complete understanding of genomic inheritance within this lineage. Assembly-based rice pangenome analyses have shown that mapping short reads directly to Nipponbare may miss divergent or non-reference sequences and have revealed extensive gene presence–absence variation among diverse accessions (Zhao et al. 2018). Population-scale studies of cultivated rice have also catalogued extensive structural and gene-content variation alongside SNPs and small indels (Wang et al. 2018). More recently, a graph-based super-pangenome constructed from long-read assemblies has improved the representation of structural and gene presence–absence variation across cultivated and wild Asian and African rice (Shang et al. 2022). Although these studies demonstrate the importance of integrating multiple classes of genomic variation, they primarily examine broad populations, domestication, and interspecific diversity rather than the transmission and reorganization of genomic variation within a documented breeding pedigree.

In this study, we generated chromosome-scale assemblies of the Hattanso landrace and four cultivars representing successive stages of the Hattan breeding lineage to investigate how the documented breeding history is reflected in their genomes. Specifically, we characterized (i) genome-wide relationships among the five Hattan-lineage accessions, (ii) variation in these relationships across chromosomes and local genomic regions, and (iii) small variants, structural variants, and gene-content differences associated with lineage-defined genomic regions. Our analyses reveal how chromosome-scale assemblies and multiple classes of genomic variation can link a documented breeding pedigree to heterogeneous inheritance patterns. The resulting assemblies and variant datasets also provide resources for comparative genomic studies of Japanese sake-brewing rice.

## Materials and Methods

### Study design

The analysis included the Nipponbare reference assembly AGIS1.0 (GCF_034140825.1) and chromosome-scale assemblies of five Hattan-lineage accessions: Hattanso, Hattan No. 10, Hattan No. 35, Hattan-nishiki No. 1, and Hattan-nishiki No. 2 (Supplementary Table S1 in Additional file 1). Unless otherwise stated, all comparative analyses were restricted to the 12 nuclear chromosomes in the AGIS1.0 coordinate system.

### Plant materials

Seed stocks of Hattanso (HGB 19950247) and Hattan No. 10 (HGB 19942166) were obtained from the Hiroshima Agricultural Gene Bank. The Hattanso stock originated from accession HO45 maintained at Kyushu University. Seeds of Hattan No. 35, Hattan-nishiki No. 1, and Hattan-nishiki No. 2 were sourced from stocks maintained at the Hiroshima Prefectural Technology Research Institute, Agricultural Technology Research Center. Seeds of all five accessions were sown in pots at the Agricultural Technology Research Center in Higashihiroshima, Hiroshima, Japan, in March 2025. Young leaves were collected from one plant per accession approximately 2–3 weeks after sowing and immediately processed for high-molecular-weight (HMW) DNA extraction.

### Genomic DNA extraction and sequencing

HMW genomic DNA was extracted using the MagAttract HMW DNA Kit (Qiagen, Hilden, Germany). DNA concentration was measured with the Qubit dsDNA HS Assay (Thermo Fisher Scientific, Waltham, MA, USA), while HMW DNA fragment size, quantity, and quality were assessed using a Femto Pulse system (Agilent Technologies, Santa Clara, CA, USA). DNA was sheared to a target fragment size of 10–15 kb using a g-TUBE (Covaris, Woburn, MA, USA) and purified with SMRTbell cleanup beads (Pacific Biosciences, Menlo Park, CA, USA).

Libraries were prepared using the SMRTbell Prep Kit 3.0 and SMRTbell Barcoded Adapter Plate 3.0 (Pacific Biosciences). Sequencing was performed in-house on a PacBio Sequel IIe system using SMRT Cells 8M, Sequel II Sequencing Kit 2.0, an on-plate loading concentration of 90 pM, and 30-h movies. HiFi reads were generated from filtered subreads using the CCS algorithm in SMRT Link v11.1.

Read statistics were summarized using the stats -a command in SeqKit (Shen et al. 2024), with coverage calculated relative to the final nuclear assembly span of each accession. Per-accession statistics are provided in Supplementary Table S1.

### Genome assembly and reference-guided scaffolding

For each accession, the HiFi FASTQ file was used directly as input to hifiasm v0.19.8-r603 (Cheng et al. 2021; Cheng et al. 2022). Primary contigs were generated using --primary, with default values retained for all other settings. The contigs were screened for potential contamination using FCS-GX v0.5.5 (Astashyn et al. 2024), after which organelle-like and rDNA-dominated contigs were identified and removed with cleanup_short_contigs (https://github.com/nigyta/cleanup_short_contigs). The cleaned primary contigs were scaffolded against AGIS1.0 (GCF_034140825.1) with RagTag v2.1.0 (Alonge et al. 2022). The AGIS1.0 mitochondrial (NC_011033.1), plastid (NC_001320.1), and mitochondrial plasmid B1 (NC_001751.1) records were excluded as scaffolding targets. RagTag was run with its default gap setting, inserting 100-bp gaps between joined contigs. The resulting scaffold sequences were used in downstream analyses.

### Genome assembly quality assessment

Assembly completeness was assessed for all six genomes using BUSCO v6.0.0 (Tegenfeldt et al. 2025) in genome mode with the poales_odb12 dataset (6,282 orthologs). The five Hattan assemblies were also assessed using aggregate consensus quality values (QVs) estimated from HiFi reads with Merqury v1.3 (Rhie et al. 2020).

### Gene-model construction and representative proteomes

AGIS1.0/RAP-DB-supported protein-coding gene models were transferred to each scaffolded assembly using LiftOn v1.0.9 (Chao et al. 2025; Shumate and Salzberg 2021). Partial mappings were excluded, coding-sequence-aware polishing was enabled, and additional gene copies were recovered at a minimum sequence identity of 0.99. De novo gene models were predicted using Helixer v0.3.6 (Holst et al. 2026) with the land_plant model, and non-overlapping predictions were added using AGAT v1.7.0 (Dainat 2026). Helixer-derived transcripts encoding proteins shorter than 50 amino acids were removed, whereas transferred and reference-derived models were retained. Coding and protein sequences were extracted with gffread v0.12.8 (Pertea and Pertea 2020), and the longest protein isoform per gene was selected to construct the representative proteome.

Annotation quality was assessed using BUSCO v6.0.0 with the poales_odb12 dataset and OMArk v0.4.1 with the Oryza lineage (Manni et al. 2021; Nevers et al. 2025; Tegenfeldt et al. 2025). Detailed settings are provided in Supplementary Methods (Additional file 2).

### Pangenome graph construction and extraction of graph-derived single-nucleotide variants (SNVs)

A pangenome graph was constructed from the AGIS1.0 reference and five chromosome-scale Hattan assemblies using Minigraph-Cactus v3.1.2 (Hickey et al. 2024), with AGIS1.0 as the reference. The resulting graph, Variant Call Format (VCF), and Hierarchical Alignment format (HAL) were used in downstream analyses.

The primary SNV dataset was extracted from the six-genome Cactus pangenome analysis. Variants in repetitive regions were excluded, and only complete, homozygous, biallelic SNV sites across AGIS1.0 and all five Hattan genomes were retained.

### HiFi read mapping and independent SNV calling

PacBio HiFi reads from each of the five accessions were aligned to AGIS1.0 using pbmm2 v1.17.0 (PacificBiosciences/pbmm2), a PacBio-specific wrapper for minimap2 (Li 2018) with the --preset CCS option. The resulting Binary Alignment/Map format (BAM) files served as common inputs for independent small-variant calling.

SNVs were independently called using DeepVariant v1.9.0 (Poplin et al. 2018; Yun et al. 2020) and Longshot v1.0.0 (Edge and Bansal 2019). DeepVariant was run using the google/deepvariant:1.9.0 container with the PACBIO model, whereas Longshot was run with the parameters -P 0.1 -F -A -D 10:500:50 and an explicit sample identifier. Concordant records were extracted using BCFtools v1.22 (Danecek et al. 2021) with the bcftools isec -n=2 -w1 command. Subsequent filtering for complete-case genotypes and site type yielded the DeepVariant–Longshot (DVLS) validation alignment.

### Genome-wide and local phylogenetic analyses

The Cactus-derived SNV alignment was used for the primary phylogenetic analysis, whereas the independently generated DVLS alignment provided cross-pipeline validation. To assess protein-sequence support for the SNV-based whole-genome topology, a concatenated alignment of representative proteins comprising 11,056,173 amino acid residues was generated using OrthoFinder v3.1.5 (Emms et al. 2026).

Maximum-likelihood phylogenies were inferred using IQ-TREE v3.1.3 (Wong et al. 2026). ModelFinder with ascertainment-bias correction was applied to the SNV alignments, with TVM+F+ASC selected for the primary Cactus analysis (Kalyaanamoorthy et al. 2017). Branch support was evaluated using 1,000 replicates each for Shimodaira–Hasegawa approximate likelihood ratio test (SH-aLRT) (Guindon et al. 2010) and ultrafast bootstrap (Hoang et al. 2018). Pairwise maximum-likelihood distances were estimated under the same fitted model. The trees were interpreted as unrooted; AGIS1.0 was positioned at the base for visualization only. Chromosome-specific trees were evaluated across all 12 chromosomes, and local phylogenies were inferred from 1-Mb windows on Chr8 and Chr12. Cactus–DVLS concordance was assessed in the focal regions. Detailed procedures are provided in Supplementary Methods.

### Genome-wide inheritance-mosaic classification and refinement of candidate transition intervals

AGIS1.0 positions aligned across all six genomes were extracted from the Cactus HAL alignment using HAL utilities. Positions overlapping the merged repeat mask were excluded, leaving 244,499,272 callable bp across the 12 nuclear chromosomes. Complete-case Cactus SNVs were summarized in non-overlapping 100-kb, 500-kb, and 1-Mb windows. The 500-kb windows defined the primary inheritance mosaic; the 100-kb and 1-Mb windows were used to refine transitions and assess broad-scale support, respectively.

Within each window, SNVs were summarized by allele-sharing pattern among the Hattan accessions, using AGIS1.0 as the reference. Windows were assigned to lineage-event or accession-specific categories defined in Supplementary Table S4 according to the dominant allele-sharing pattern. Assignment required three criteria: a callable fraction of ≥0.20, at least three informative SNVs, and a matching dominant pattern with a fraction of ≥0.60. Assignments with a callable fraction of ≥0.40 and at least five informative SNVs were considered high confidence. Other qualifying windows were classified as provisional, whereas windows not meeting the assignment criteria remained unresolved. Adjacent windows with the same assignment were merged without bridging unresolved gaps.

Candidate transitions were refined to intervals bounded by the nearest differently assigned 100-kb windows. These intervals represent changes between adjacent window-based mosaic classifications. DVLS validation required exact matches for chromosome, position, reference allele, and alternate allele. Missing genotypes were not treated as reference. Detailed classification criteria are provided in Supplementary Methods.

### Comparison of Shuho SNV states with Hattan No. 10 and Hattan No. 35 using public sequence data

Publicly available 100-bp paired-end reads for Shuho (DRR054268; Yano et al. 2016) were mapped to the six-genome pangenome graph with vg giraffe v1.66.0 (Sirén et al. 2021). Allelic support for Hattan No. 10 and Hattan No. 35 was compared at 25,833 biallelic SNVs distinguishing the two cultivars, with the Hattan No. 35 allele designated as the pedigree-consistent state. Allele-constrained genotype likelihoods were used to compare support for the two cultivar states. Regional support was summarized in 500-kb windows containing at least five covered markers at which the Hattan No. 35 allele was nonreference relative to AGIS1.0.

Sensitivity analyses examined the potential effects of reference-allele status and read depth and compared graph-based and linear mapping, using BWA-MEM2 v2.2.1 (Vasimuddin et al. 2019) for linear mapping. Detailed calling, filtering, and sensitivity-analysis procedures are provided in Supplementary Methods.

### Functional annotation of SNVs and small indels

Complete, non-conflicting, biallelic SNVs and simple 1–50-bp insertions or deletions were extracted from the Cactus VCF. Callable variants were normalized against AGIS1.0 and annotated using a custom AGIS1.0 database in snpEff v5.4c (Cingolani et al. 2012). The analysis was restricted to variants within lineage-assigned regions whose genotype partitions matched the corresponding regional categories, retaining only the most severe predicted consequence for each variant. Predicted HIGH-impact variants were evaluated for reference identity, callability, gene-model quality, assembly context, and exact-allele DVLS support. To account for nearby variation, callable variants within 10 bp were incorporated by reconstructing and reannotating accession-specific homozygous compound haplotypes. Variants were retained only if the HIGH-impact consequence persisted and no unresolved technical or gene-model concerns remained. Detailed procedures are provided in Supplementary Methods.

### Structural-variant evidence integration

Structural variants (SVs) comprised insertions, deletions, and inversions of >50 bp, together with translocation/breakend events. SVs were standardized to AGIS1.0 coordinates and integrated across three evidence families: (i) long-read consensus calls from Sniffles2 v2.7.0 (Smolka et al. 2024), cuteSV v2.1.2 (Jiang et al. 2020), and pbsv v2.11.0 (https://github.com/PacificBiosciences/pbsv); (ii) assembly-based evidence from whole-genome alignments generated with nucmer (MUMmer4 v4.0.1) (Marçais et al. 2018) and interpreted with SyRI v1.7.1 (Goel et al. 2019); and (iii) graph-based evidence comprising large insertion and deletion alleles from the Cactus graph. Events were matched within and across Hattan accessions and annotated for gene, repeat, projected query-gap, and inheritance-mosaic context. The primary multi-evidence set required support from at least two of these evidence families and had to pass the structural-annotation filters. Replicated single-family events were retained separately. Detection patterns summarized observed calls across the five Hattan accessions. Detailed integration criteria are provided in Supplementary Methods.

### Pangene construction and gene presence–absence variation (PAV) curation

A modified get_pangenes workflow (Contreras-Moreira et al. 2023) grouped corresponding loci into pangenes across AGIS1.0 and the five Hattan assemblies using whole-genome alignments and gene coordinates. Assignments were reconciled with OrthoFinder protein orthogroups, and presence–absence patterns specific to breeding steps or accessions were selected as initial PAV candidates. Candidates identified in Hattan assemblies were projected onto AGIS1.0 coordinates using the Cactus HAL alignment. Candidate evaluation considered local DNA and protein correspondence, surrounding gene order, gene models, SVs, and HiFi reads. Supported PAVs required resolved local presence–absence states supported by assembly sequences and HiFi reads. Local absence with a highly similar copy elsewhere was classified as positional PAV. Genes affected by the same structural event were counted as one independent locus. Detailed criteria are provided in Supplementary Methods.

### Transposable-element annotation and spatial relationships with SVs, PAV loci, and candidate transition intervals

Transposable elements (TEs) were annotated with HiTE v3.3.2 (Hu et al. 2024). We examined whether curated SVs, primary lineage-associated PAV loci, and candidate transition intervals occurred within or near TEs. Detailed procedures are provided in Supplementary Methods.

### Functional annotation of six rice proteomes and enrichment analysis of lineage- and variant-associated gene sets

The AGIS1.0 and five Hattan proteomes were functionally annotated using EnTAP v2.3.0 (Hart et al. 2020), and Gene Ontology (GO) terms (Ashburner et al. 2000; The Gene Ontology Consortium 2026) and Kyoto Encyclopedia of Genes and Genomes (KEGG) identifiers (Kanehisa et al. 2025) were standardized across all six proteomes. Enrichment targets included genes in lineage-assigned intervals, genes carrying final primary HIGH-impact variants, genes overlapped by SVs supported by at least two evidence families, and primary lineage-associated PAV gene sets. Each curated pangene unit was counted once, with the corresponding functionally annotated units used as the background. GO and pathway enrichment were evaluated using one-sided hypergeometric tests followed by Benjamini–Hochberg correction (Benjamini and Hochberg 1995). Only tests with at least ten annotated target units were performed, and significance was defined at false discovery rate (FDR) ≤ 0.05. Detailed annotation models, minimum target-size requirements, pathway filters, GO–pathway comparison criteria, and sensitivity analyses are described in Supplementary Methods.

### Use of generative AI in manuscript preparation

OpenAI Codex was used to assist with English-language editing, manuscript organization, consistency checking across the manuscript and supplementary materials, and preparation or revision of analysis and visualization code. All scientific content, analyses, interpretations, references, and generated outputs were reviewed and verified by the authors, who take full responsibility for the content of the manuscript.

## Results

### Genome assembly and annotation quality across Hattan-lineage genomes

We analyzed the Hattanso landrace and four derived cultivars representing successive stages of the documented Hattan breeding lineage (Fig. 1A). The chromosome-scale assemblies ranged from 375.78 to 378.94 Mb in length, with scaffold N50 values of 30.25–30.43 Mb and 99.8% BUSCO completeness, and Merqury quality values ranged from 65.80 to 67.05 (Fig. 1B; Supplementary Table S1).

**Figure 1.**
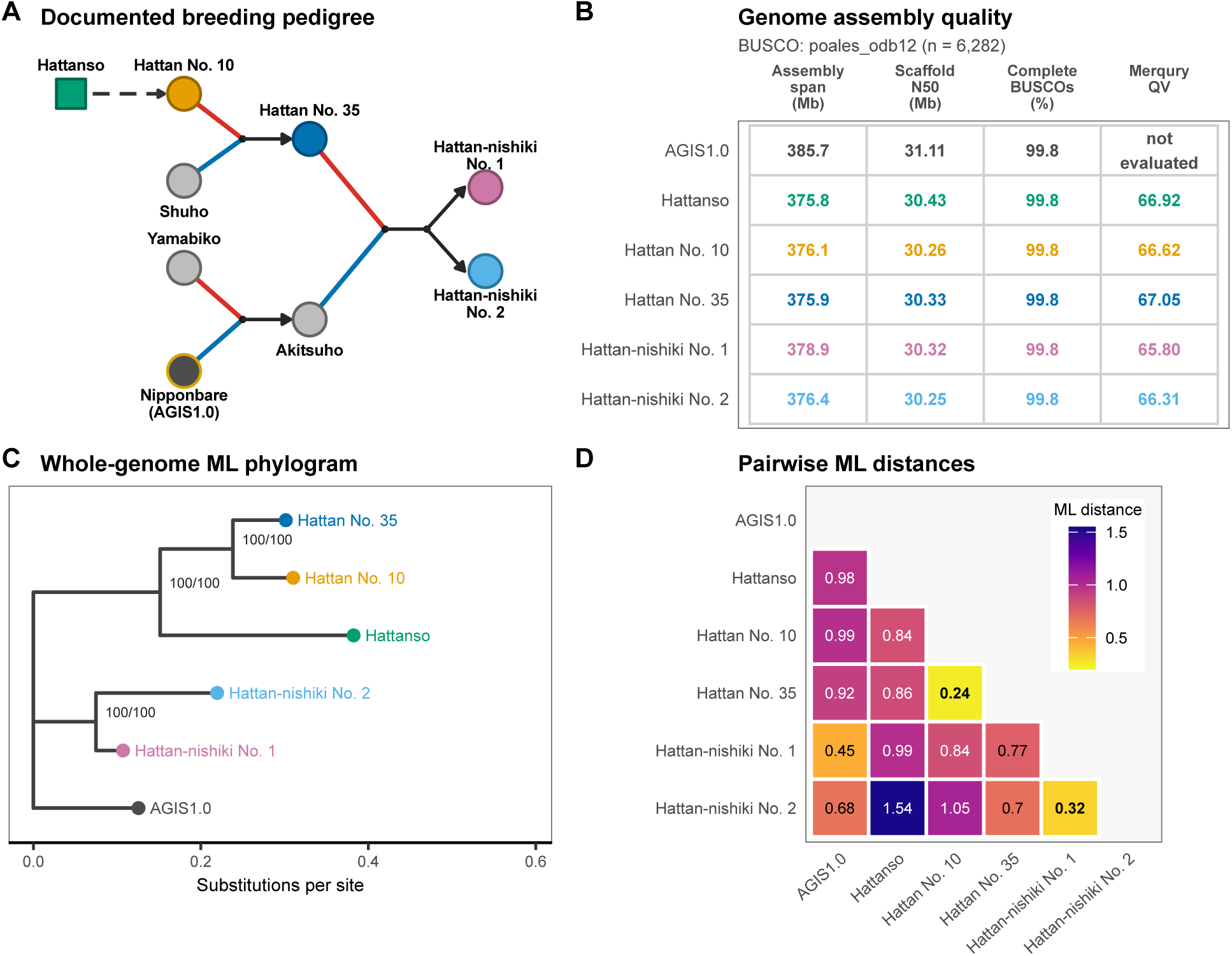
Breeding pedigree, genome assembly quality, and whole-genome relationships of AGIS1.0 and five Hattan-lineage genomes. (A) Simplified documented pedigree of the focal Hattan breeding series and the Nipponbare contribution through Akitsuho. The square denotes the Hattanso landrace, circles denote cultivars, the dashed arrow denotes pure-line selection, and red and blue segments identify maternal and paternal parents, respectively. Chromosome-scale assemblies were generated for the five color-coded Hattan accessions. Grey circles indicate pedigree materials without chromosome-scale assemblies; publicly available Shuho reads were analyzed separately. AGIS1.0 is the Nipponbare reference assembly. (B) Assembly span, scaffold N50, complete BUSCO percentage, and aggregate Merqury QV. BUSCO v6.0.0 was run using the poales_odb12 dataset (6,282 orthologs). A directly comparable read-based Merqury QV was not evaluated for AGIS1.0. (C) Maximum-likelihood phylogram inferred from 174,754 Cactus graph-derived biallelic SNVs under TVM+F+ASC. Internal labels indicate SH-aLRT and ultrafast-bootstrap support, each based on 1,000 replicates. AGIS1.0 is displayed on the basal branch for visualization and was not treated as a biological outgroup. (D) Lower-triangle matrix of maximum-likelihood genetic distances estimated from the same Cactus SNV set. The Hattan No. 10–Hattan No. 35 and Hattan-nishiki No. 1–Hattan-nishiki No. 2 comparisons are shown in bold.

Protein-annotation quality was consistent across AGIS1.0 and the five Hattan-lineage genomes. Representative proteomes were 99.7% complete according to protein-mode BUSCO analysis. OMArk classified 98.53–98.59% of conserved hierarchical orthologous groups as single or duplicated and 93.40–93.66% of gene models as taxonomically consistent, with no likely contaminants detected (Supplementary Table S2).

### Genome-wide relationships among Hattan-lineage accessions

Genome-wide relationships among the five Hattan-lineage accessions and AGIS1.0 were evaluated using two SNV datasets: 174,754 homozygous biallelic SNVs with non-missing genotypes across all six genomes, obtained from the Cactus graph-based analysis, and 168,542 independently called SNVs from DeepVariant plus Longshot (DVLS). Both datasets identified Hattan No. 10 and Hattan No. 35 as the closest pair, followed by Hattan-nishiki Nos. 1 and 2 (Fig. 1C, D and Supplementary Fig. S1 in Additional file 3). Maximum-likelihood distances were strongly correlated across all 15 sample pairs (Pearson’s r = 0.959; Supplementary Fig. S1 and Supplementary Table S3).

Both SNV datasets recovered the same principal unrooted topology. Hattan No. 10 and Hattan No. 35 formed a clade grouped with Hattanso, whereas Hattan-nishiki Nos. 1 and 2 formed a second clade. The three principal internal branches received 100% support from both SH-aLRT and ultrafast bootstrapping in both analyses. The 11,056,173-residue concatenated OrthoFinder protein alignment recovered the same topology, supporting its reproducibility across variant-calling pipelines and sequence representations (Fig. 1C and Supplementary Fig. S1).

### Heterogeneity in genomic relationships across chromosomes and local regions

Chromosome-level phylogenies revealed variation in genomic relationships across the 12 nuclear chromosomes. The Hattan No. 10–Hattan No. 35 clade was recovered on 9 of 12 chromosomes, although ultrafast-bootstrap support ranged from 50% to 100%. The complete whole-genome topology was recovered only on Chr1, Chr9, and Chr11 (Supplementary Fig. S2 in Additional file 4).

Window-based analyses of Chr8 and Chr12 revealed further variation within individual chromosomes (Supplementary Fig. S3 in Additional file 5). Of the 17 non-overlapping windows of up to 1 Mb on Chr8 that contained at least 50 parsimony-informative sites, two recovered the complete whole-genome topology, two contained the Hattan No. 10–Hattan No. 35 topology feature, and 11 contained a local Hattan No. 35–Hattan-nishiki No. 2 feature. Among eight windows on Chr12 meeting the same criterion, the corresponding counts were zero, one, and four.

The regional patterns were consistent between the Cactus and DVLS datasets, with genotype patterns agreeing at 91.00–99.56% of exactly overlapping variant sites across three focal regions. Refinement using 100-kb windows identified discontinuous candidate blocks totaling 2.90 Mb with a Hattan No. 35–Hattan-nishiki No. 2 shared pattern and 3.70 Mb with a Hattan-nishiki No. 2-private pattern.

### Window-based classification of the Hattan genomic mosaic

To characterize this local heterogeneity, we classified 776 non-overlapping 500-kb windows based on the dominant allele-sharing patterns among the five Hattan accessions (Fig. 2A). Of these, 402 were assigned to lineage-pattern categories covering 200.2 Mb (51.91%) of the nuclear genome. The largest category comprised windows dominated by SNVs at which the four derived cultivars shared an allele different from that of the sampled Hattanso accession. These windows covered 20.2% of the AGIS1.0 nuclear reference sequence. Other major categories included a nonreference state shared across the Hattan lineage (9.3%) and the Hattan No. 35–Hattan-nishiki transition pattern (8.2%) (Fig. 2B). The remaining windows were classified as unresolved, with detailed subcategories and integrated block assignments presented in Supplementary Fig. S4 in Additional file 6 and Supplementary Table S4.

**Figure 2.**
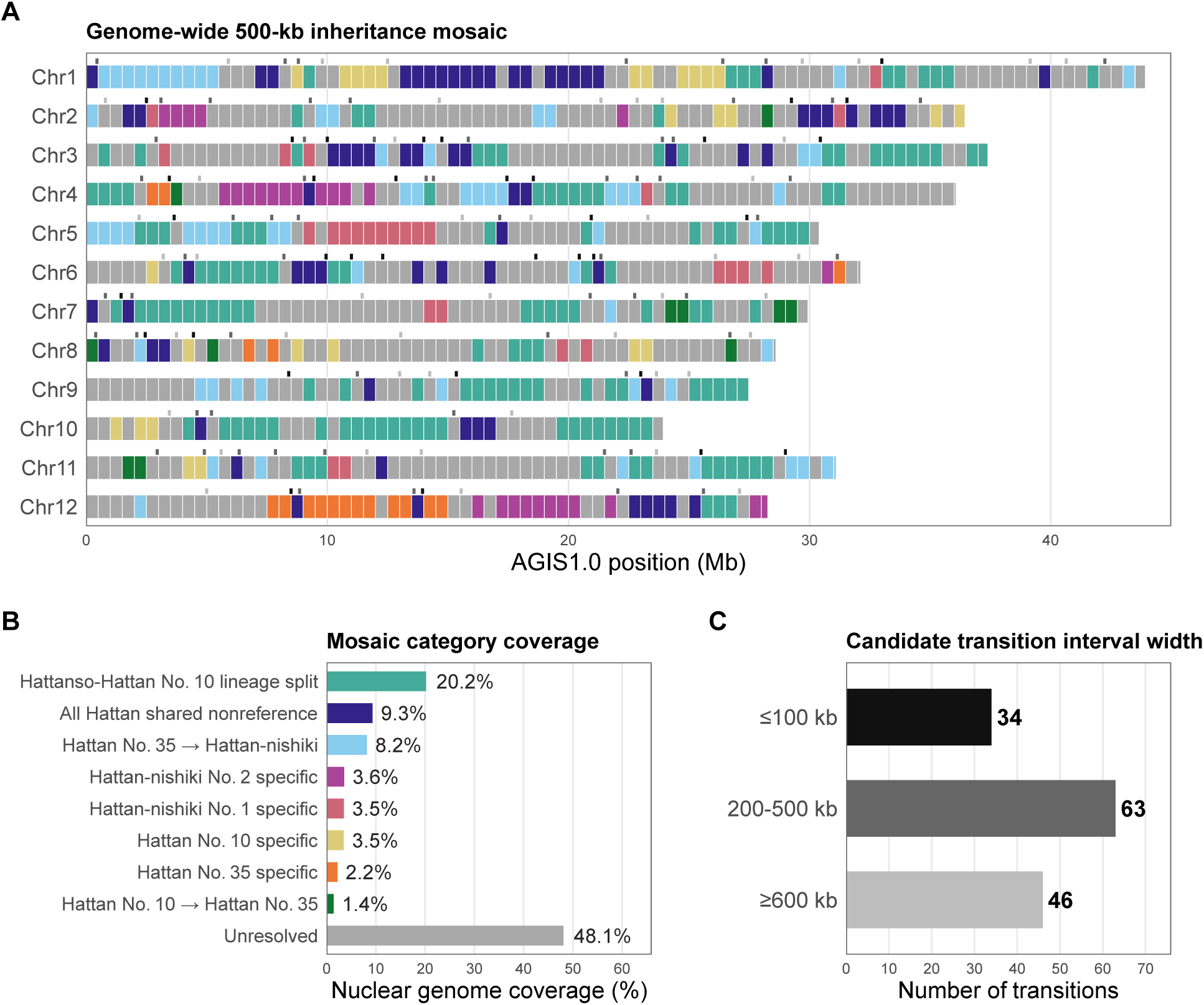
Genome-wide inheritance mosaic and candidate transition intervals. (A) Integrated 500-kb inheritance mosaic across the 12 AGIS1.0 nuclear chromosomes. Colored tiles show lineage-pattern categories inferred from Cactus graph-derived variants. The unresolved category is shown in grey. Upper ticks mark candidate transitions, with shading indicating the interval-width classes summarized in panel C. (B) Physical coverage of the nuclear genome by the assigned lineage categories and the combined unresolved category. Arrows indicate abbreviated category names associated with successive breeding transitions. The Hattanso–Hattan No. 10 lineage-split category combines orientation-complementary patterns that separate Hattanso from Hattan No. 10 and the later Hattan cultivars. Lineage-associated categories accounted for 200.2 Mb, or 51.91% of the nuclear genome. (C) Width distribution of the 143 candidate category-transition intervals. Assignment-confidence shading, detailed unresolved subcategories, and spatial DVLS validation status are shown in Supplementary Fig. S4.

We identified 143 candidate transitions between lineage-pattern categories, 34 of which were narrowed to intervals of ≤100 kb using local 100-kb classifications (Fig. 2A, C). Independent comparison with DVLS provided complementary support for the mosaic classifications (Supplementary Fig. S4A). Among 138,256 sites with exactly matching reference and alternate alleles, DVLS genotypes supported the expected alternate-allele carriers at 123,700 sites (89.47%).

Allele-level concordance with Shuho was assessed using publicly archived paired-end reads (DRR054268) at biallelic SNVs distinguishing Hattan No. 10 from Hattan No. 35. Among markers at which Hattan No. 35 carried the nonreference allele, 7,492 were covered after graph mapping and 7,240 after linear mapping. Of the informative reads at these markers, 80.88% after graph mapping and 79.82% after linear mapping supported the Hattan No. 35 allele. The Hattan No. 35 state was also favored in 64 of 66 500-kb windows containing at least five covered Hattan No. 35-nonreference markers. This widespread concordance is consistent with the documented contribution of Shuho to Hattan No. 35 (Supplementary Fig. S5 in Additional file 7 and Supplementary Table S5).

The genomic-mosaic analysis also distinguished the sister cultivars Hattan-nishiki Nos. 1 and 2, both derived from the documented Hattan No. 35 × Akitsuho cross (Figs. 2A and 3B). Cultivar-specific mosaic windows totaled 13.50 Mb (3.50% of the nuclear genome) in Hattan-nishiki No. 1 and 13.77 Mb (3.57%) in Hattan-nishiki No. 2. Although similar in total length, these states occupied different genomic regions, revealing distinct realized outcomes of the same recorded cross.

**Figure 3.**
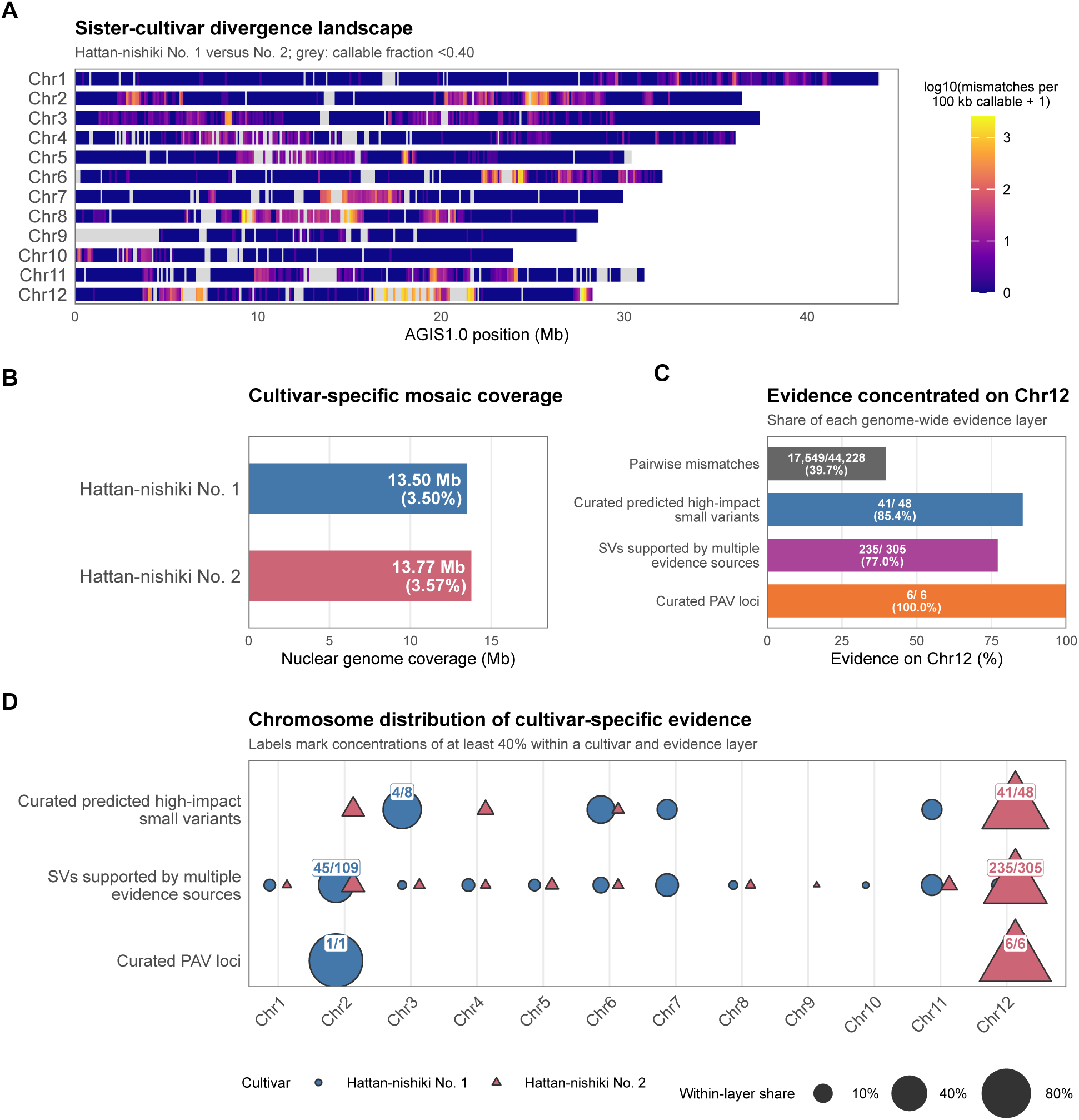
Genomic divergence between Hattan-nishiki Nos. 1 and 2. (A) Pairwise mismatch landscape normalized by callable sequence in non-overlapping 100-kb windows across the 12 AGIS1.0 chromosomes. Colors represent log₁₀ (mismatches per 100 kb of callable sequence + 1), from dark blue for lower values to yellow for higher values. Grey denotes windows with a callable fraction below 0.40. (B) Coverage of cultivar-specific categories in the integrated 500-kb inheritance mosaic. (C) Fraction of each genome-wide Hattan-nishiki No. 2-specific evidence layer located on Chr12. (D) Chromosome distribution of cultivar-specific small variants, SVs, and supported lineage-associated PAV loci for both sister cultivars. Color and shape distinguish the two cultivars, and point area represents the fraction of each cultivar-specific evidence layer assigned to a chromosome. Labels mark concentrations of at least 40%.

Differentiation between the sister cultivars was particularly pronounced on Chr12, which accounted for 17,549 of their 44,228 genome-wide mismatches (39.7%) (Fig. 3A). Small-variant and structural-variant evidence assigned to the Hattan-nishiki No. 2-specific state was similarly concentrated on Chr12, which also contained all six supported PAV loci in this category (Fig. 3C, D). This convergence across variant classes identifies Chr12 as a major focus of differentiation between the sister cultivars and illustrates the spatial heterogeneity of the Hattan genomic mosaic.

### Integrated variation landscape across the Hattan lineage

We annotated 194,618 primary-callable Cactus-derived SNVs and 1–50-bp indels. Of these, 116,426 occurred in lineage-pattern regions and exhibited allele-sharing patterns that matched the categories assigned to their respective 500-kb windows. SnpEff initially classified 428 variants as HIGH impact. Following evaluation of sequence context, compound haplotypes, gene-model quality, callability, and regional support, 385 small variants were retained in the final curated set of predicted high-impact variants (Fig. 4A, B and Supplementary Table S6).

**Figure 4.**
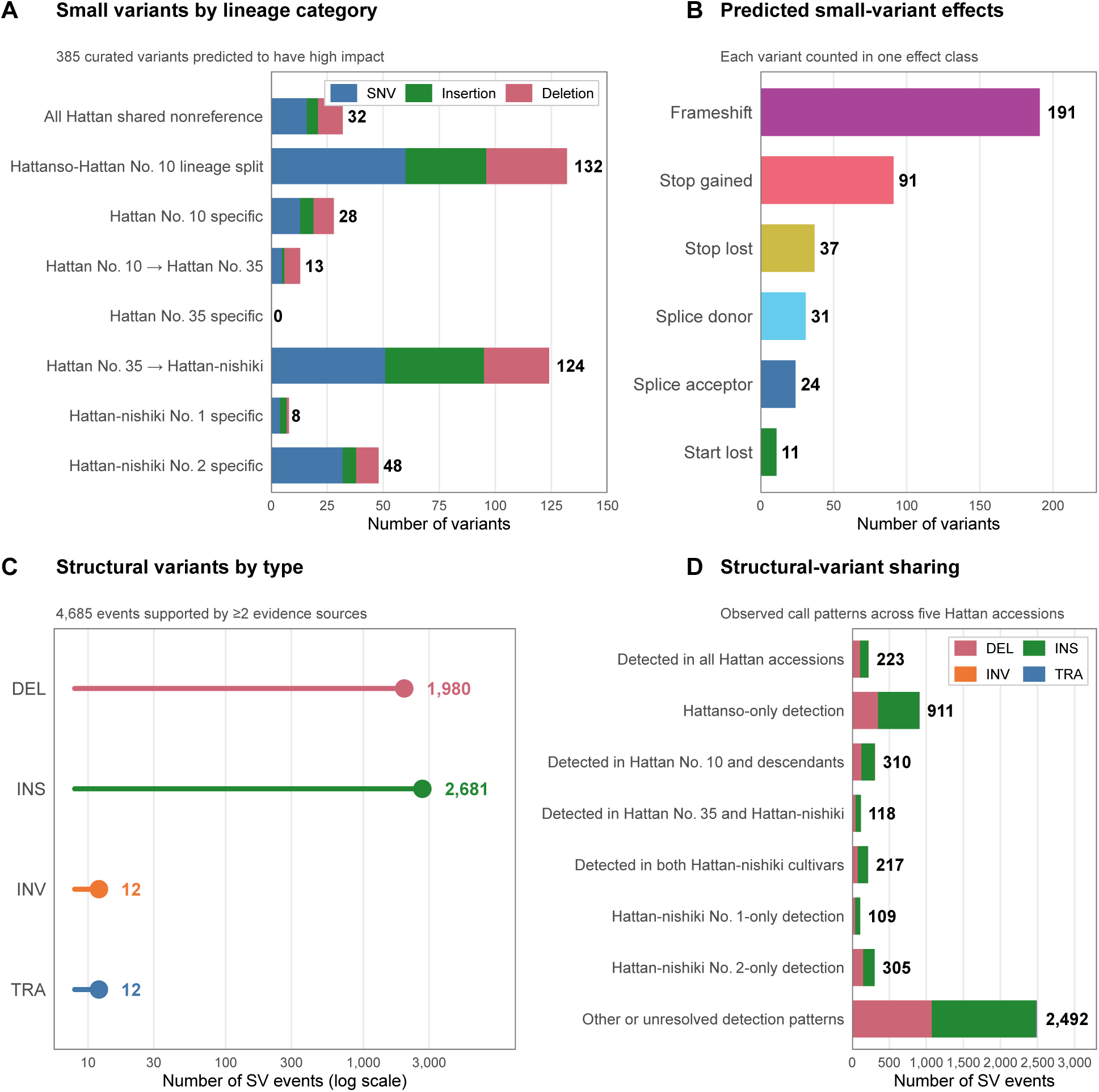
Predicted high-impact small variants and structural variants across the Hattan lineage. (A) Counts of 385 curated small variants (SNVs and 1–50-bp insertions and deletions) predicted to have high impact, grouped by lineage category. The Hattanso–Hattan No. 10 lineage-split category combines orientation-complementary patterns. (B) Predicted consequence classes of the same 385 variants. Variants assigned multiple consequence terms were counted once according to the displayed priority order. (C) Type composition of 4,685 structural-variant (SV) events >50 bp that were supported by at least two evidence families and passed structural-annotation filters. (D) Detection patterns of the same 4,685 SVs across the five Hattan accessions. The Hattanso-only category denotes events detected only in the sampled Hattanso genome. The remaining categories include non-persistent detection patterns and unresolved assignments.

Larger-scale variation was characterized by integrating three evidence sources: long-read calls, whole-genome assembly alignments, and the Cactus graph. This analysis identified 11,951 nonredundant structural-variant (SV) events, of which 4,685 were supported by multiple evidence families and passed structural-annotation filters. The multi-evidence set comprised 1,980 deletions, 2,681 insertions, 12 inversions, and 12 translocations (Fig. 4C, D and Supplementary Table S7). Of these 4,685 events, 2,193 showed lineage-compatible detection patterns. Among these 2,193 events, 512 overlapped annotated genes, and 335 also agreed with the corresponding mosaic classifications (Fig. 5D).

**Figure 5.**
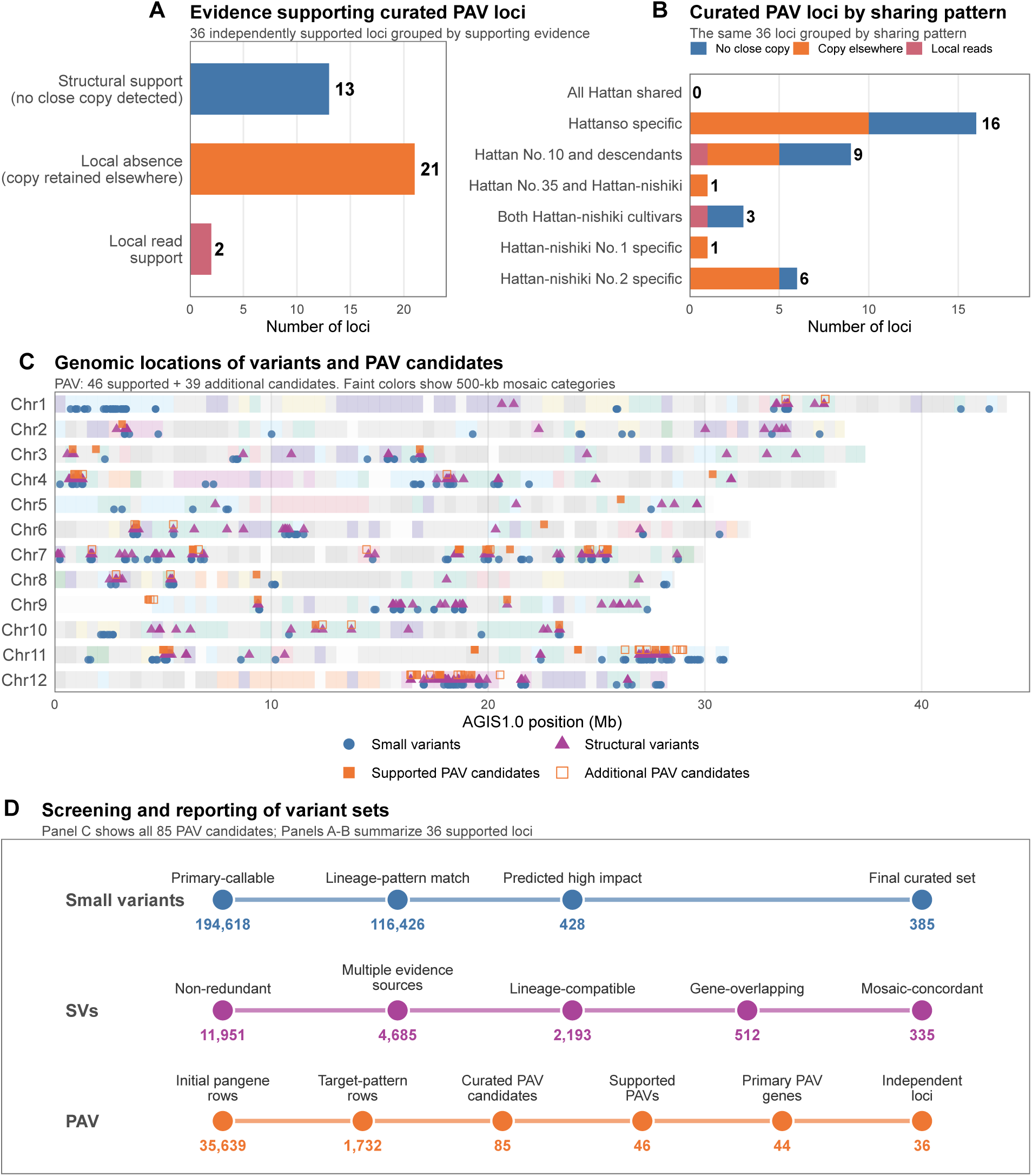
Candidate gene presence–absence variation and genomic locations of variant sets across the Hattan lineage. Of the 85 PAV candidates, the primary supported lineage-associated set comprised 44 genes at 36 independent loci. (A) Evidence and copy-context classes for this set. Direct structural support refers to evidence of gene-deleting SVs from multiple evidence families, without a detected close full-length copy elsewhere. Positional absence denotes local absence with a close copy retained elsewhere and can also have direct SV support. The remaining supported loci rely on local sequence and read evidence. (B) Distribution of the same 36 loci among lineage-associated sharing patterns. Colors indicate the classes in panel A. (C) AGIS1.0 chromosomal locations of 385 curated predicted high-impact small variants, 335 lineage-compatible, gene-overlapping, mosaic-concordant SVs, and all 85 PAV candidate units. Filled and open squares indicate the 46 candidates with assembly-sequence and HiFi read support and the 39 requiring further evaluation, respectively. Candidate positions were projected onto AGIS1.0 coordinates. Faint background colors show the 500-kb mosaic categories. (D) Screening and reporting stages for the three variant classes. The PAV path starts with 35,639 initial pangene rows and 1,732 rows matching breeding-step or accession-specific patterns, followed by the 85 curated candidate units plotted in panel C. Subsequent stages show 46 supported gene-level PAVs and the primary lineage-associated set of 44 genes, which were grouped into 36 independent loci for panels A and B. Supplementary Table S9 lists all 85 candidate units with supporting observations.

We next evaluated the spatial distribution of multi-evidence SVs relative to classified TEs, considering insertions and deletions separately. Compared with chromosome-wise randomized positions, both SV types more frequently had at least one boundary within 1 kb of a classified TE (Supplementary Fig. S6A, C, D in Additional file 8). Deletions also more frequently had a boundary within 100 bp of a TE and were enriched for direct TE overlap. In contrast, insertions showed no detectable enrichment within 100 bp and were depleted for direct TE overlap. TE-proximity patterns differed among TE classes (Supplementary Fig. S6B and Supplementary Table S8).

We next evaluated lineage-associated differences in gene content. Locus-level evaluation retained 85 PAV candidates, of which 46 had local sequence and HiFi read support (Supplementary Table S9). Of these, 44 exhibited breeding-step-or accession-specific presence–absence patterns, representing 36 independent loci (Fig. 5A, B). Candidates requiring further evaluation included partial sequences and cases with unresolved locus correspondence, gene structure, or read support.

The chromosome-aligned distributions of the 385 small variants, 335 SVs, and 85 PAV candidates provide an integrated view of nucleotide, structural, and gene-content variation across the Hattan genomes (Fig. 5C). We also reviewed 15 focal SV loci for structural evidence, agreement with inheritance patterns, gene overlap and model quality, TE context, and functional evidence (Supplementary Table S10).

Functional enrichment analysis further characterized genes from the lineage-associated regions. The Hattan No. 10-specific set showed the most consistent signal, with pectinesterase-related GO functions converging with the Pentose and glucuronate interconversions pathway across annotation-sensitivity analyses. The Hattan-nishiki No. 2-specific set showed enrichment for diterpenoid-and terpenoid-related functions and the Diterpenoid biosynthesis pathway (Supplementary Fig. S7 in Additional file 9 and Supplementary Table S11).

### Genomic regions and alleles shared across the Hattan lineage

We identified 12,205 SNVs for which all five Hattan accessions shared the same nonreference allele. Shared-SNV density per callable megabase was highest on Chr4. Within the 1-Mb region spanning Chr4:17.5–18.5 Mb, 2,960 of 2,976 SNVs, 340 of 342 small indels, and 32 of 36 multi-evidence SVs were shared by all five accessions. Of these shared SVs, 20 had support from multiple evidence families in all five accessions (Supplementary Fig. S8 in Additional file 10 and Supplementary Table S12). Detailed procedures are provided in Supplementary Methods.

## Discussion

By integrating genome-wide and local analyses of chromosome-scale assemblies from the Hattanso landrace and four derived cultivars, we found that genomic relationships varied across chromosomes and local regions. Although Hattan No. 10 and Hattan No. 35 were the closest pair at the whole-genome scale, some chromosomes and local regions supported different groupings, including local clustering of Hattan No. 35 with Hattan-nishiki No. 2. Classification of 500-kb windows organized this heterogeneity into lineage-associated genomic states, while analyses of small variants, structural variants, and gene presence–absence variation revealed differences at the nucleotide, structural, and gene-content levels. Together, these findings show that the Hattan genomes form a mosaic in which strong genome-wide relationships coexist with distinct local patterns of allele sharing.

This genomic-mosaic interpretation extends previous studies of Japanese rice breeding, which showed that documented pedigrees can be resolved into chromosome-scale haplotype blocks and that recombination and selection reshape their genomic distribution (Yamamoto et al. 2010; Yonemaru et al. 2012). Using chromosome-scale assemblies, we examined how SNV-based relationships varied from whole genomes to individual chromosomes and local windows within the documented Hattan lineage. The regional patterns were supported by both assembly-derived and read-based SNV datasets. Public Shuho reads also broadly supported the Hattan No. 35 state at discriminating markers, consistent with the recorded Hattan No. 10 × Shuho parentage. These findings demonstrate how multiscale genomic analysis can complement pedigree records by revealing spatial heterogeneity that whole-genome relationships alone may obscure.

The contrast between Hattan-nishiki Nos. 1 and 2 illustrates the genomic mosaic revealed by this multiscale analysis. Although both cultivars were selected from the same recorded Hattan No. 35 × Akitsuho cross, segregation and recombination can produce different combinations of parental chromosome segments (Huang et al. 2009; Kwon et al. 2025).

Consistent with this expectation, the cultivars had similar total spans of cultivar-specific mosaic states, but these states occupied different genomic regions. Differentiation was particularly concentrated on Chr12, which accounted for 39.7% of their genome-wide mismatches. This chromosome also contained most of the curated small and structural variants assigned to the Hattan-nishiki No. 2-specific state, along with all six PAV loci in that category. These convergent patterns identify Chr12 as a major region of differentiation and illustrate how descendants of the same recorded cross can exhibit distinct genomic configurations.

The concentration of multiple variant classes on Chr12 indicates that local differentiation in the Hattan lineage involved nucleotide changes as well as structural and gene-content variation. SNV-defined haplotype blocks can trace allele sharing along a pedigree, whereas assembly-resolved insertions, deletions, and PAVs can reveal gene disruptions, local gene presence or absence, and nonreference sequences that may be poorly represented in an SNV-only analysis based on a single reference genome. This broader perspective is consistent with rice population and pangenome studies identifying structural and gene-content variation as substantial components of intraspecific diversity (Wang et al. 2018; Shang et al. 2022). The integrated map included 385 predicted high-impact small variants, 335 multi-evidence SVs that overlapped genes and matched mosaic classifications, and 85 PAV candidates (Fig. 5C, D). Together, these variant classes provide additional structural and gene-content resolution beyond SNP-based pedigree analysis and identify candidates for functional investigation.

Genomic features shared by Hattanso and the four derived cultivars may provide clues to the genetic background retained within the Hattan lineage. The Chr4:17.5–18.5 Mb region is particularly noteworthy because most detected variants were shared across all five accessions. This region includes wall-associated kinase genes, such as OsWAK43 and OsWAK44, previously proposed as candidates for spikelet fertility under heat stress (Lafarge et al. 2017). Phenotypic comparisons with other rice lineages would help determine whether Hattan accessions maintain spikelet fertility at high temperatures and evaluate their potential for heat-tolerance breeding.

Enrichment analysis identified biological processes potentially associated with reported Hattan traits. In the Hattan No. 10-specific set, pectinesterase-related GO terms and the Pentose and glucuronate interconversions pathway pointed to pectin modification and galacturonate metabolism. Previous work found no significant differences in measured starch and protein properties among Hattan-lineage materials (Tamaki et al. 2005b), although Hattan No. 10 exhibited relatively high grain hardness and one of the highest broken-grain rates during polishing (Tamaki et al. 2005a). Because pectin methylesterases modify plant cell walls, this signal suggests that cell-wall remodeling may contribute to the cultivar’s polishing characteristics (Jeong et al. 2015). The Hattan-nishiki No. 2-specific set showed enrichment for diterpenoid-related functions. This cultivar was bred for cultivation at higher elevations, where low temperatures constrain rice production (Tamaki et al. 2005a; Ye et al. 2009). Because rice diterpenoid phytoalexin pathways participate in biotic and abiotic stress responses (Yamamura et al. 2015; Huang et al. 2024), this enrichment may indicate functions relevant to the target cultivation environment. Grain histology, cell-wall analysis, pathway-gene expression, metabolite profiling, and relevant phenotyping would provide direct routes for testing these hypotheses.

Resolving the ancestry underlying the Hattan genomic mosaic will require additional parental and population genomic resources. Chromosome-scale assemblies of Shuho and Akitsuho from material linked to the relevant breeding stocks would enable local segments to be assigned to parental sources and extend the present SNV-level Shuho comparison to the inheritance of structural and gene-content variation. Broader sampling of Hattanso would help distinguish characteristics of the sampled accession from genetic diversity within the historical landrace population, as rice landraces can retain substantial within-cultivar variation (Olufowote et al. 1997; Yamasaki and Ideta 2013). These resources would improve the interpretation of candidate transition intervals, which currently represent changes in classified genomic states rather than confirmed recombination breakpoints. Parental assemblies and denser local marker data could help establish which intervals represent transitions between parental haplotypes and refine their locations. Incorporating these resources into the analytical framework could ultimately translate documented pedigrees into genome-resolved ancestry maps and clarify how nucleotide, structural, and gene-content variation is transmitted and reorganized during breeding.

## Conclusions

Chromosome-scale comparison of the Hattanso landrace and four derived cultivars revealed a spatially heterogeneous genomic mosaic reflecting their documented breeding relationships. Although Hattan No. 10 and Hattan No. 35 were the closest pair at the whole-genome scale, lineage-associated genomic states varied across chromosomes and local regions. Hattan-nishiki Nos. 1 and 2, sister cultivars derived from the same recorded cross, displayed distinct genomic configurations, with differentiation particularly concentrated on Chr12. Integrating nucleotide, structural, and gene-content variation across genomic scales complements pedigree records and helps prioritize candidate regions and genes for future functional and breeding studies.

## List of abbreviations

BAM: Binary Alignment/Map
BUSCO: Benchmarking Universal Single-Copy Orthologs
CCS: Circular consensus sequencing
CDS: coding sequence
DVLS: DeepVariant-plus-Longshot
FDR: false discovery rate
GO: Gene Ontology
HAL: hierarchical alignment HMW: high molecular weight
KEGG: Kyoto Encyclopedia of Genes and Genomes
LINE: Long interspersed nuclear element
LTR: Long terminal repeat
PAV: presence–absence variation
QV: quality value
SH-aLRT: Shimodaira–Hasegawa approximate likelihood-ratio test
SINE: Short interspersed nuclear element
SNP: Single-nucleotide polymorphism
SNV: single-nucleotide variant
SV: structural variant
TE: transposable element
TIR: Terminal inverted repeat
VCF: Variant Call Format.

## Declarations

### Ethics approval and consent to participate

Not applicable. This study used rice plants and publicly available sequence data and did not involve human participants or animals.

### Consent for publication

Not applicable.

### Availability of data and materials

The PacBio HiFi reads generated for the five Hattan-lineage accessions have been deposited in the DDBJ Sequence Read Archive under BioProject PRJDB45902 and run accessions DRR1213530–DRR1213534. The corresponding chromosome-scale genome assemblies have been deposited in DDBJ under sequence accession numbers AP054507–AP054588. Accession numbers for each Hattan-lineage accession are provided in Supplementary Table S1. The public Shuho reads analyzed in this study are available from the DDBJ Sequence Read Archive under accession DRR054268. Supporting data are provided in the supplementary information files.

Genome annotations and comparative genomic data, including selected variant callsets and detailed PAV evaluation records, are available in Figshare (https://doi.org/10.6084/m9.figshare.35048491). The analysis and figure-generation code are available in a separate Figshare record (https://doi.org/10.6084/m9.figshare.35048533).

### Authors’ contributions

RN performed the bioinformatic analyses from raw sequence data onward and wrote the original draft. ZK and AF cultivated the plants. MK collected plant samples, extracted DNA, prepared sequencing libraries, and performed sequencing. SY prepared the data visualizations. SM conceived the study, cultivated the plants, and reviewed and edited the manuscript. HB conceived the study and reviewed and edited the manuscript. All authors read and approved the final manuscript.

## Acknowledgements

We thank all members of the Laboratory of Genome Informatics at Hiroshima University for their cooperation and valuable support.

## Supplementary figure legends

**Supplementary Figure S1. Whole-genome relationship reproducibility across sequence representations.** (A–C) Maximum-likelihood topologies inferred from 174,754 Cactus graph-derived SNVs, 168,542 independently called DeepVariant-plus-Longshot SNVs, and the 11,056,173-residue concatenated OrthoFinder representative-protein alignment, respectively. Internal labels indicate SH-aLRT/ultrafast-bootstrap support. The trees are interpreted as unrooted, with AGIS1.0 displayed at the base for visualization. (D) Pairwise maximum-likelihood distance comparison for the two SNV datasets across all 15 sample pairs.

**Supplementary Figure S2. Chromosome-level phylogenies across the Hattan lineage.** Maximum-likelihood trees were inferred independently for the 12 AGIS1.0 nuclear chromosomes from Cactus-derived SNVs under the fixed TVM+F+ASC model. Node labels give SH-aLRT/ultrafast-bootstrap support. The trees are interpreted as unrooted, with AGIS1.0 displayed at the base for visualization. Panel titles report the number of SNV sites and indicate whether the complete whole-genome topology or the Hattan No. 10–Hattan No. 35 clade was recovered.

**Supplementary Figure S3. Genomic relationships differ among non-overlapping windows of up to 1 Mb on Chr8 and Chr12.** The chromosomes were partitioned into non-overlapping 1-Mb windows, with terminal partial windows retained. A maximum-likelihood tree was inferred for each window containing at least 50 parsimony-informative sites, yielding 17 analyzed windows on Chr8 and eight on Chr12. The three topology features were evaluated independently. A window matching the complete whole-genome topology also contains the Hattan No. 10–Hattan No. 35 clade and therefore appears as both green and blue. Orange tiles indicate an alternative local Hattan No. 35–Hattan-nishiki No. 2 clade. White tiles indicate analyzed windows in which the displayed feature was not recovered. Grey intervals contained fewer than 50 parsimony-informative SNV sites and were not analyzed.

**Supplementary Figure S4. Detailed inheritance-mosaic categories, DeepVariant-plus-Longshot support, and candidate transition intervals.** (A) The genome-wide 500-kb inheritance mosaic with unresolved subclasses shown separately. Main tiles show the detailed Cactus-derived lineage-pattern categories. Lighter shades indicate provisional assignments. Narrow lower tracks summarize independent validation based on exact allele matches using the DeepVariant-plus-Longshot dataset. Black solid lines indicate replication of the expected allele pattern, black points indicate mixed or discordant support, and light-grey solid lines indicate insufficient or unresolved evidence. Upper ticks indicate candidate transition-interval width. (B) Physical genome coverage of the assigned lineage categories and the four unresolved subclasses. Assigned categories follow the order used in Figure 2. Unresolved subclasses are grouped below from pattern-based ambiguity to data limitation. (C) Width classification of the 143 candidate transition intervals. The intervals indicate changes in classified genomic state rather than demonstrated molecular recombination breakpoints.

**Supplementary Figure S5. Shuho support for the Hattan No. 35 allele across mapping and filtering conditions.** The same set of 7,736,854 publicly archived 100-bp reads from DRR054268 were evaluated after graph mapping with vg giraffe and linear mapping to the 12 AGIS1.0 nuclear chromosomes with BWA-MEM2. (A) Percentages of mapped and properly paired reads. Points are used because the vertical axis is restricted to the observed high-mapping range. (B) Fractions of informative reads supporting the Hattan No. 35 allele across all Hattan No. 10/Hattan No. 35-discriminating markers and the subsets in which the Hattan No. 35 allele was reference or nonreference relative to AGIS1.0. (C) Support for the nonreference Hattan No. 35 allele after requiring one to five informative reads per marker. (D) Fractions of non-tied 500-kb windows favoring Hattan No. 35 after requiring three, five, or ten covered nonreference markers per window. A window favored Hattan No. 35 when the sum of its marker-level log10 likelihood ratios was positive. Labels show the principal five-marker result: 64 of 66 windows after graph mapping and 63 of 65 after linear mapping.

**Supplementary Figure S6. Transposable-element context of structural variation.** (A) Observed-to-null ratios for direct overlap with classified AGIS1.0 transposable elements and proximity within 100 bp or 1 kb of these elements (TEs) among 4,685 structural variants (SVs) >50 bp that were supported by at least two evidence families and passed structural-annotation filters. Results are shown for all SVs and separately for 2,681 insertions and 1,980 deletions. The horizontal line at 1 indicates the null expectation. (B) Observed-to-null ratios for SV proximity within 1 kb of DNA terminal-inverted-repeat transposons (DNA TIR), Helitrons, long interspersed nuclear elements (LINEs), long terminal repeat retrotransposons (LTRs), and short interspersed nuclear elements (SINEs). Values above 1 indicate enrichment and values below 1 indicate depletion relative to chromosome-wise randomization. (C, D) Cumulative distributions of the distance from the nearest SV boundary to the nearest classified TE for all 2,681 insertions (C) and all 1,980 deletions (D). Solid lines show the observed distributions. Dashed lines and grey bands show the mean and 95% interval, respectively, from 3,000 chromosome-wise circular permutations. Vertical dotted lines mark 100 bp and 1 kb. A distance of 0 indicates that a boundary overlapped or directly abutted a classified TE. Complete TE-overlap, proximity, and sensitivity results are provided in Supplementary Table S8.

**Supplementary Figure S7. Functional enrichment across lineage-associated gene sets.** FDR-adjusted GO terms and KEGG pathways are shown for genes in 500-kb mosaic windows assigned to the Hattan No. 10-specific and Hattan-nishiki No. 2-specific categories. The horizontal axis shows −log10 of the FDR-adjusted P value, and point size denotes fold enrichment. Green points indicate GO terms and blue points indicate KEGG pathways. Circles indicate results significant in at least two annotation models. Primary and sensitivity results, including target sets below the minimum annotated target size, are provided in Supplementary Table S11.

**Supplementary Figure S8. Shared nonreference variation across the Hattan lineage and the Chr4 focal region.** (A) Density of SNVs carrying the same nonreference allele in all five Hattan accessions, shown in 500-kb windows across the 12 AGIS1.0 nuclear chromosomes. Density is expressed per callable megabase; colors use a log10 (density + 1) scale. Grey windows have a callable fraction below 0.40. The outlined interval marks Chr4:17.5–18.5 Mb. (B) Counts of SNVs, small insertions and deletions, and multi-evidence SVs in 100-kb windows across Chr4:16.5–19.5 Mb. Grey bars show all variants, blue bars show variants shared by all five accessions, and narrow dark bars show shared SVs with multi-family support in each accession. SVs are assigned by their left reference anchor. The lower tracks show callable fractions and the positions of *OsWAK43* and *OsWAK44*; the dashed line marks a callable fraction of 0.40. Shading marks Chr4:17.5–18.5 Mb. (C) Percentage of all variants that are shared by the five accessions, calculated separately within the focal interval and in the rest of the nuclear genome. Labels show shared/all variant counts and the corresponding percentage. SNVs and small indels use the complete-case callable Cactus dataset, whereas SVs use the multi-evidence event set. Shared SVs represent matched event detections across accessions. Coordinates refer to AGIS1.0, and counts are descriptive summaries of linked variation. Supporting data are provided in Supplementary Table S12.

## Additional files

**Additional file 1 (.xlsx).** Supplementary Tables S1–S13. Sample and assembly metadata, comparative analyses, and variant and enrichment results.

**Additional file 2 (.docx).** Supplementary Methods. Detailed analytical procedures, classification criteria, and sensitivity analyses.

**Additional file 3 (.tif).** Supplementary Figure S1. Reproducibility of genome-wide relationships across sequence representations.

**Additional file 4 (.tif).** Supplementary Figure S2. Chromosome-level phylogenies across the Hattan lineage.

**Additional file 5 (.tif).** Supplementary Figure S3. Phylogenetic relationships in focal windows on chromosomes 8 and 12.

**Additional file 6 (.tif).** Supplementary Figure S4. Detailed mosaic categories and validation against an independent SNV dataset.

**Additional file 7 (.tif).** Supplementary Figure S5. Allelic support from publicly archived Shuho reads.

**Additional file 8 (.tif).** Supplementary Figure S6. Transposable-element context of structural variation.

**Additional file 9 (.tif).** Supplementary Figure S7. Functional enrichment of lineage-and variant-associated gene sets.

**Additional file 10 (.tif).** Supplementary Figure S8. Shared nonreference variation and the chromosome 4 focal region.

