## Supplemental Figures for "Chromosome-scale comparative genomics reveals a genomic mosaic across the documented Hattan sake-brewing rice lineage"

#### Whole-genome relationship reproducibility across sequence representations

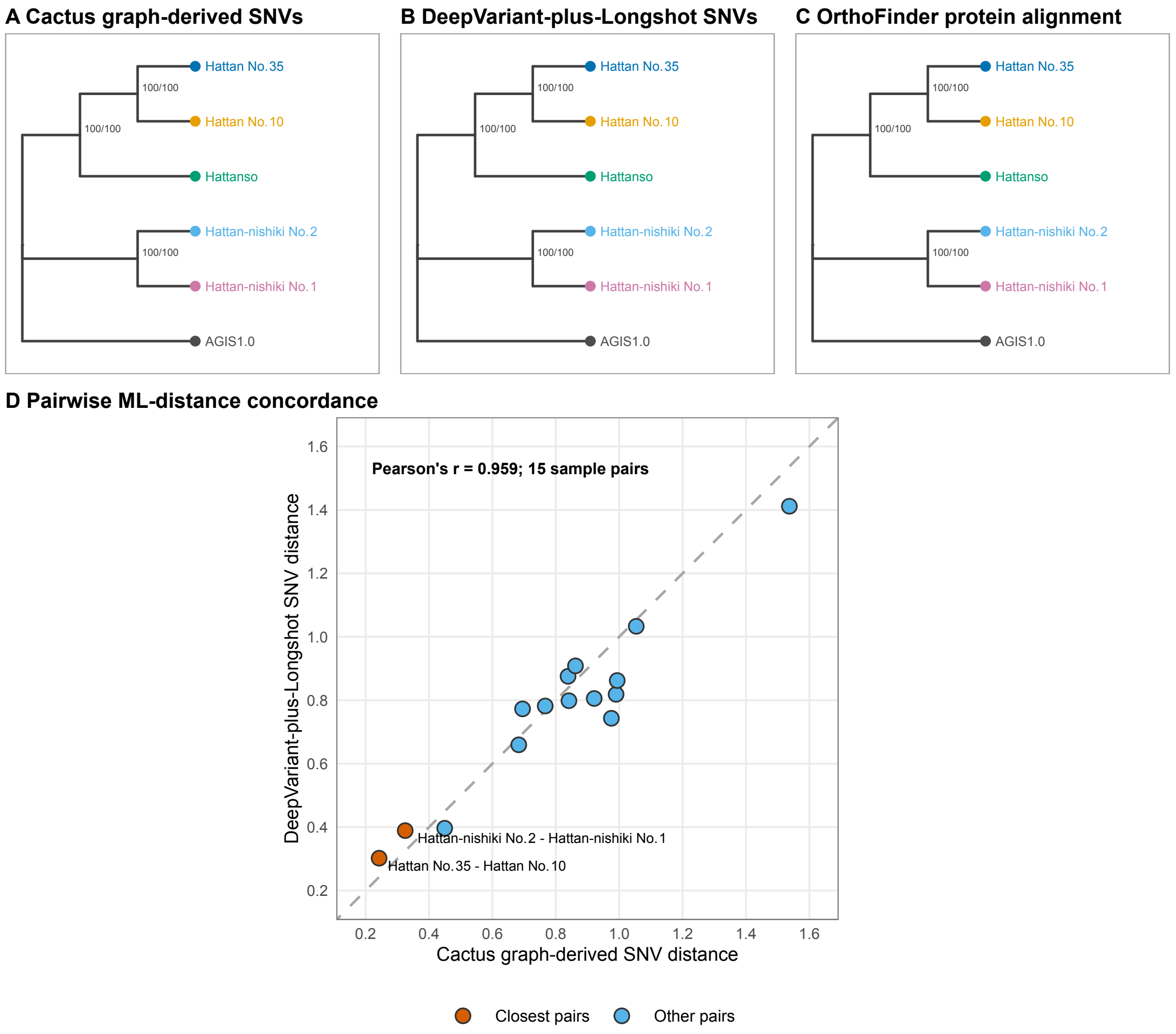

### Chromosome-level phylogenies of AGIS1.0 and five Hattan-lineage genomes

Cactus-derived SNVs | fixed TVM+F+ASC model | node labels: SH-aLRT/ultrafast bootstrap (%)

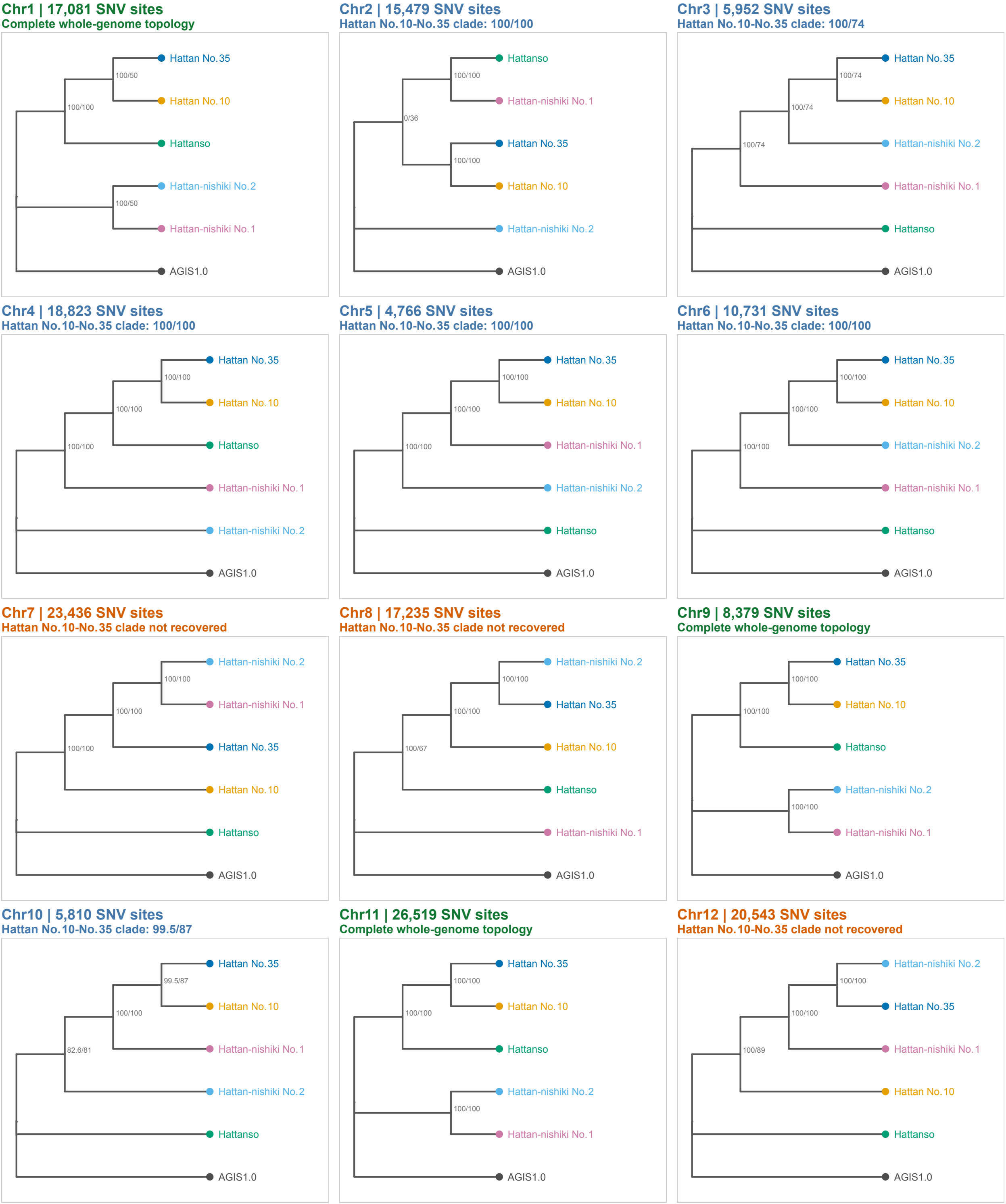

#### Genomic relationships differ among 1-Mb windows on Chr8 and Chr12

A maximum-likelihood tree was inferred separately for each analyzed 1-Mb window.
The three features below were scored independently.

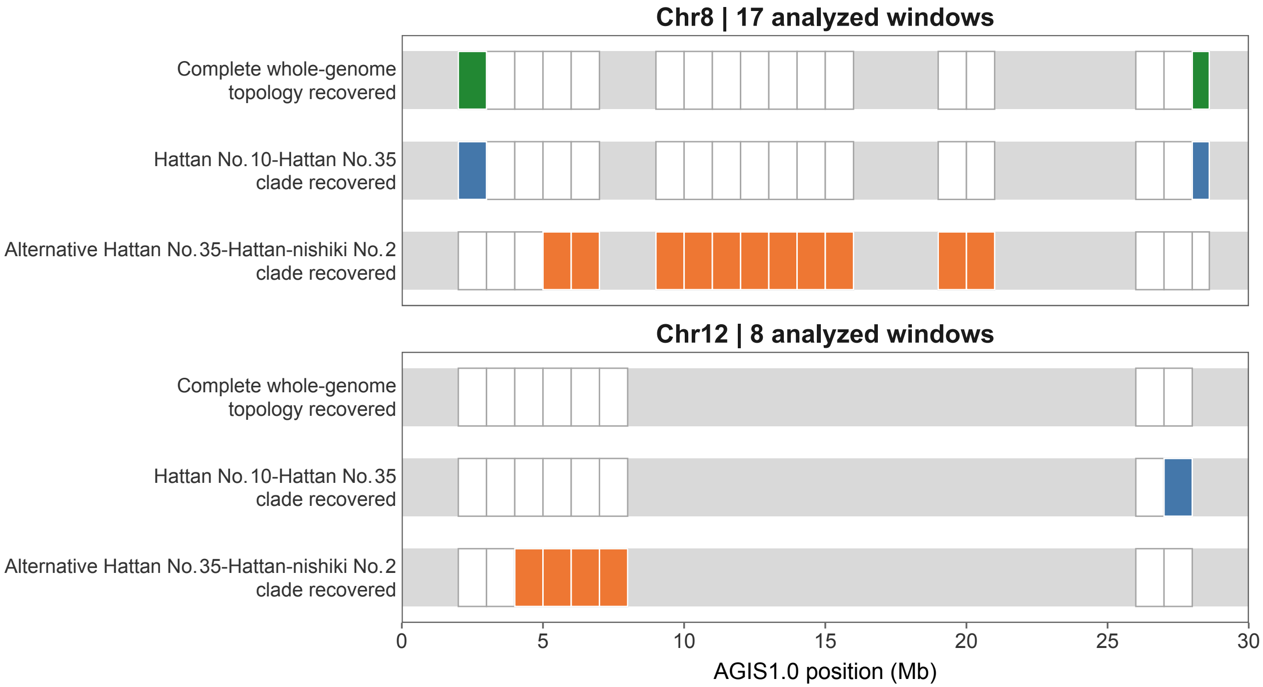

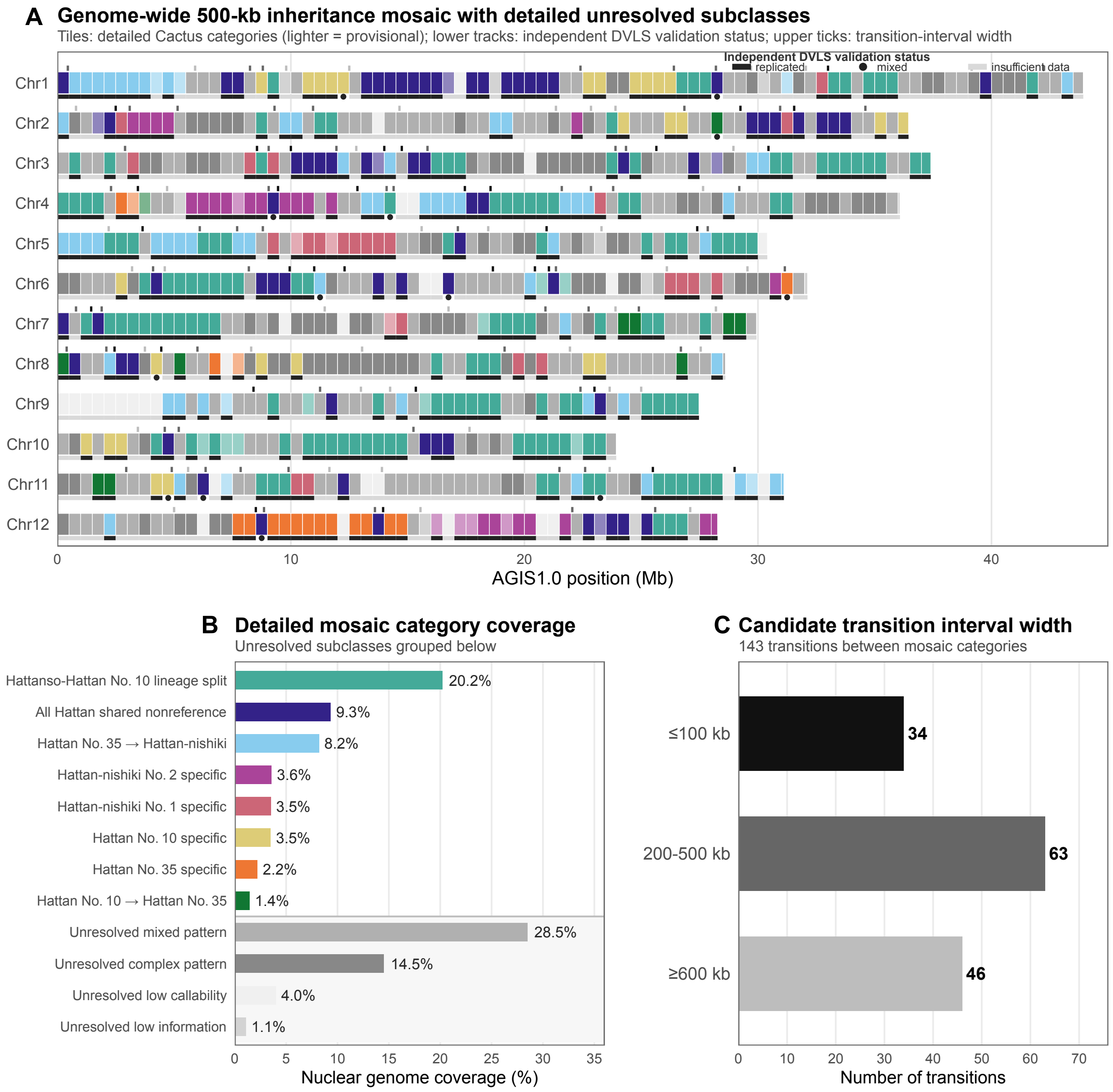
A
Genome-wide 500-kb inheritance mosaic with detailed unresolved subclasses
Tiles: detailed Cactus categories (lighter = provisional); lower tracks: independent DVLS validation status; upper ticks: transition-interval width
Independent DVLS validation status
replicated
mixed
insufficient data
Chr1
Chr2
Chr3
Chr4
Chr5
Chr6
Chr7
Chr8
Chr9
Chr10
Chr11
Chr12
0
10
20
30
40
AGIS1.0 position (Mb)
B
Detailed mosaic category coverage
Unresolved subclasses grouped below
Hattanso-Hattan No. 10 lineage split
20.2%
All Hattan shared nonreference
9.3%
Hattan No. 35 → Hattan-nishiki
8.2%
Hattan-nishiki No. 2 specific
3.6%
Hattan-nishiki No. 1 specific
3.5%
Hattan No. 10 specific
3.5%
Hattan No. 35 specific
2.2%
Hattan No. 10 → Hattan No. 35
1.4%
Unresolved mixed pattern
28.5%
Unresolved complex pattern
14.5%
Unresolved low callability
4.0%
Unresolved low information
1.1%
0
5
10
15
20
25
30
35
Nuclear genome coverage (%)
C
Candidate transition interval width
143 transitions between mosaic categories
≤100 kb
34
200-500 kb
63
≥600 kb
46
0
10
20
30
40
50
60
70
Number of transitions

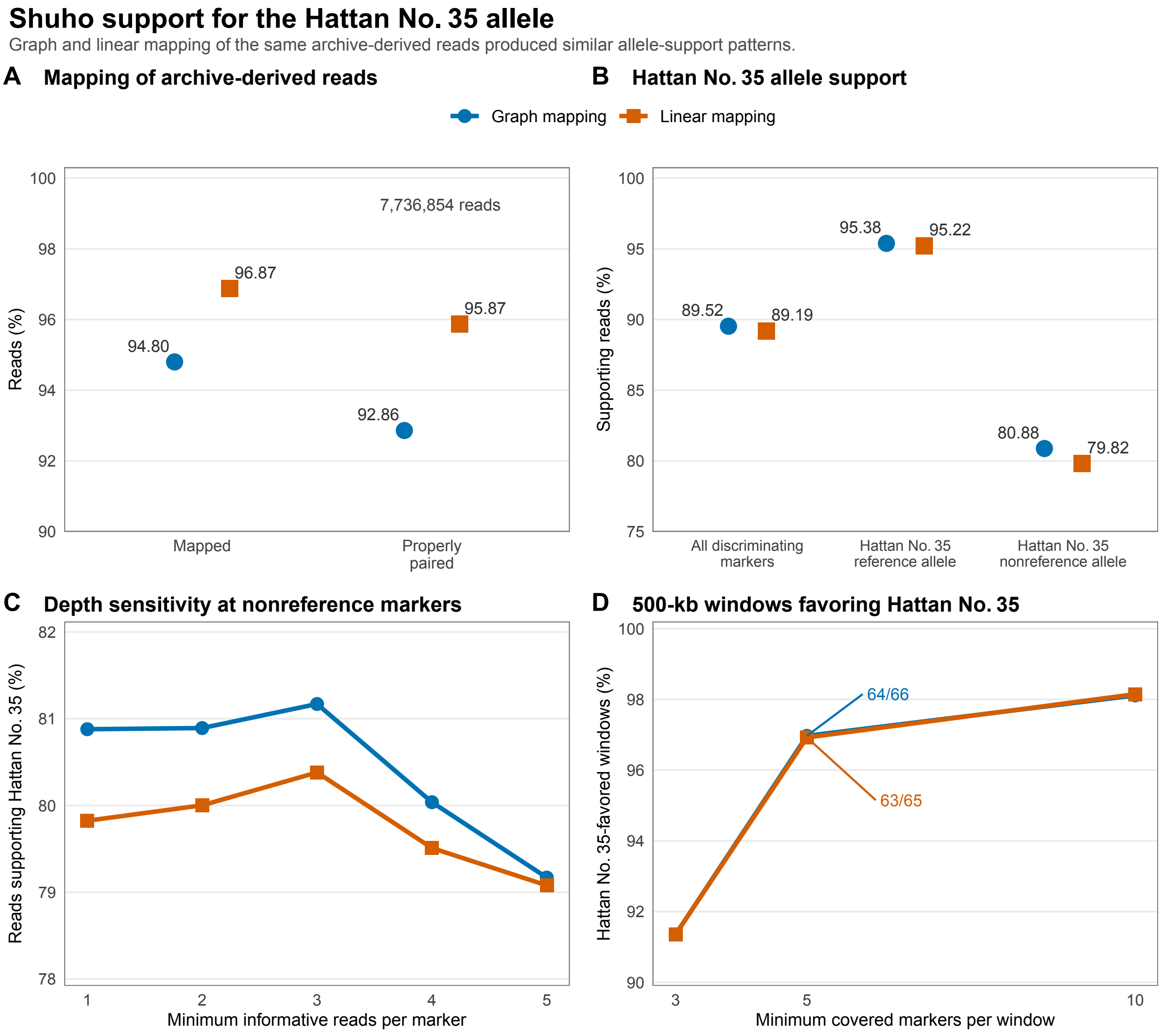
Shuho support for the Hattan No. 35 allele
Graph and linear mapping of the same archive-derived reads produced similar allele-support patterns.
A Mapping of archive-derived reads
B Hattan No. 35 allele support
Graph mapping
Linear mapping
Reads (%)
7,736,854 reads
96.87
94.80
95.87
92.86
Mapped
Properly paired
Supporting reads (%)
95.38
95.22
89.52
89.19
80.88
79.82
All discriminating markers
Hattan No. 35 reference allele
Hattan No. 35 nonreference allele
C Depth sensitivity at nonreference markers
Reads supporting Hattan No. 35 (%)
Minimum informative reads per marker
D 500-kb windows favoring Hattan No. 35
Hattan No. 35-favored windows (%)
64/66
63/65
Minimum covered markers per window

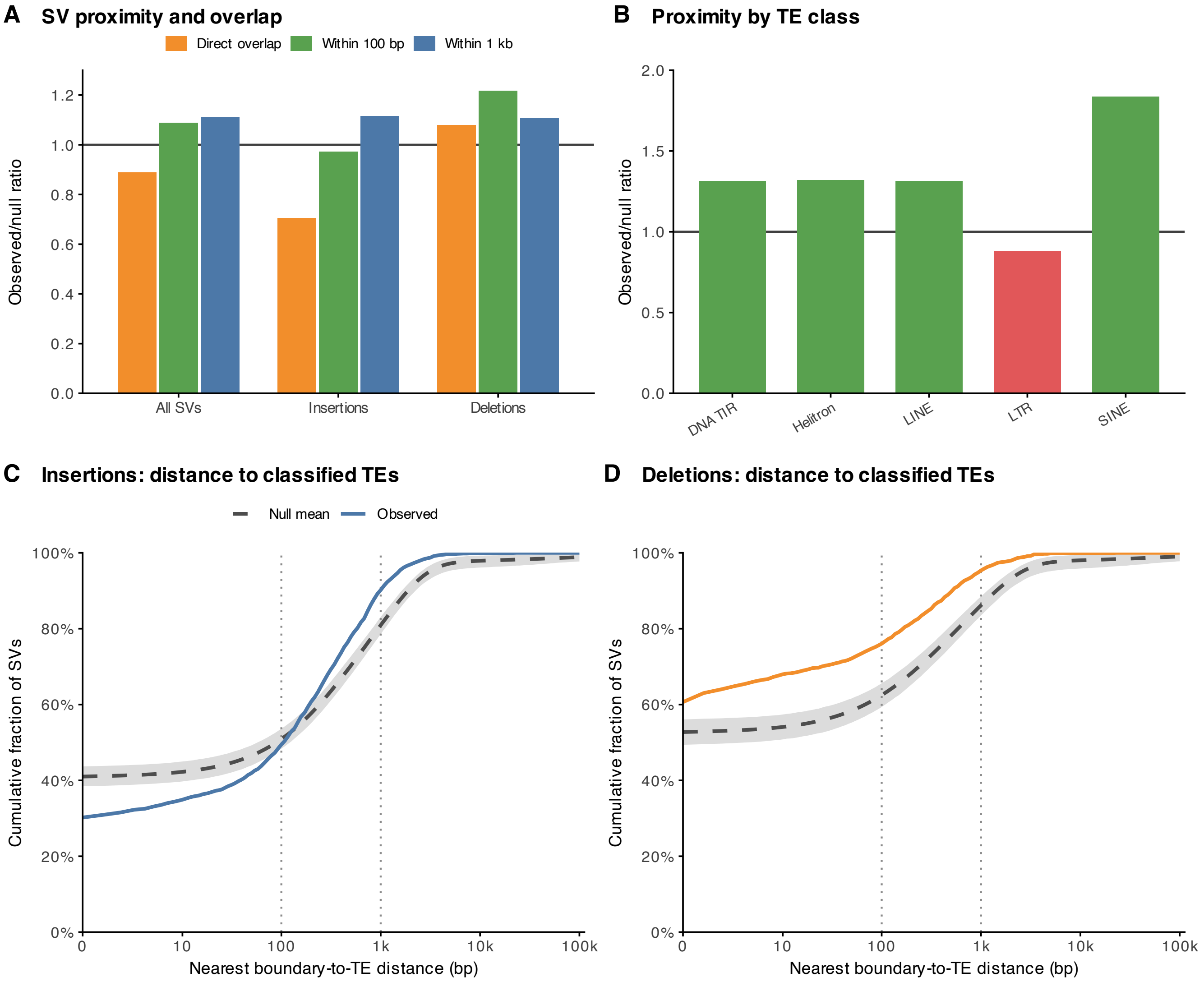
A
SV proximity and overlap
Direct overlap
Within 100 bp
Within 1 kb
Observed/null ratio
0.0
0.2
0.4
0.6
0.8
1.0
1.2
All SVs
Insertions
Deletions
B
Proximity by TE class
Observed/null ratio
0.0
0.5
1.0
1.5
2.0
DNA TIR
Helitron
LINE
LTR
SINE
C
Insertions: distance to classified TEs
Null mean
Observed
Cumulative fraction of SVs
0%
20%
40%
60%
80%
100%
0
10
100
1k
10k
100k
Nearest boundary-to-TE distance (bp)
D
Deletions: distance to classified TEs
Cumulative fraction of SVs
0%
20%
40%
60%
80%
100%
0
10
100
1k
10k
100k
Nearest boundary-to-TE distance (bp)

#### Functional enrichment across lineage-associated gene sets

Green points show GO terms and blue points show KEGG pathways. Full results are in Supplementary Table S11.

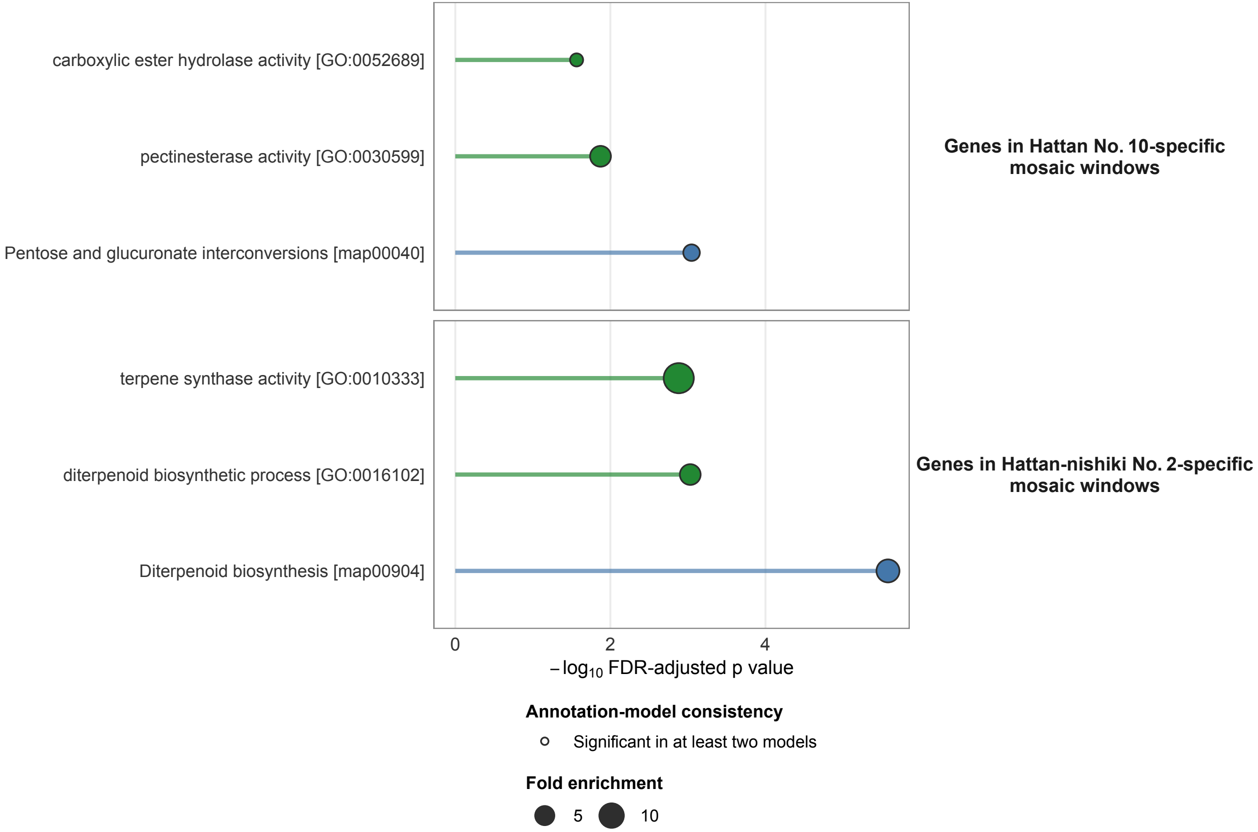

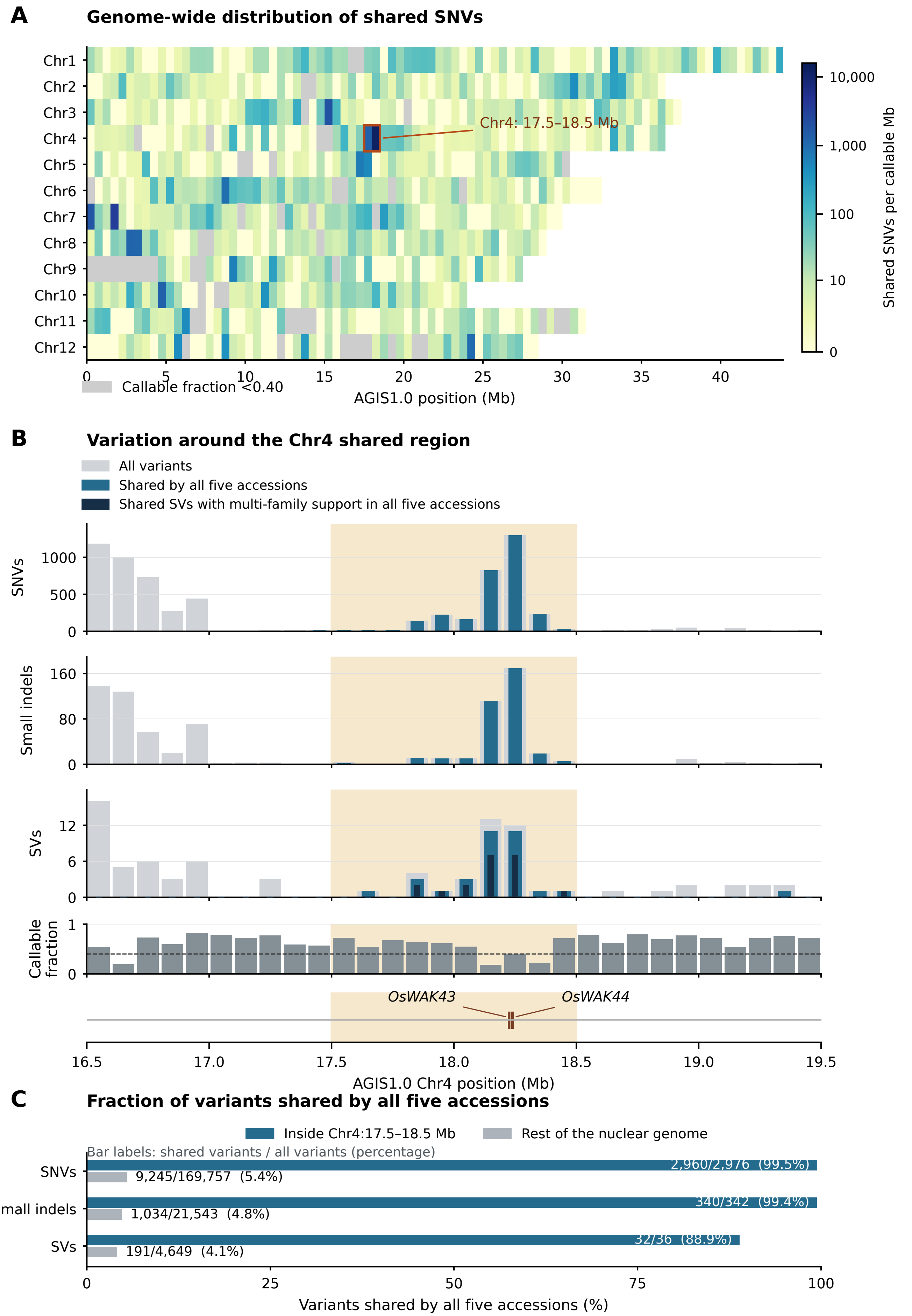

A
Genome-wide distribution of shared SNVs
Chr1
Chr2
Chr3
Chr4
Chr5
Chr6
Chr7
Chr8
Chr9
Chr10
Chr11
Chr12
Chr4: 17.5–18.5 Mb
Callable fraction <0.40
AGIS1.0 position (Mb)
Shared SNVs per callable Mb
10,000
1,000
100
10
0
B
Variation around the Chr4 shared region
All variants
Shared by all five accessions
Shared SVs with multi-family support in all five accessions
SNVs
Small indels
SVs
Callable fraction
OsWAK43
OsWAK44
AGIS1.0 Chr4 position (Mb)
C
Fraction of variants shared by all five accessions
Inside Chr4:17.5–18.5 Mb
Rest of the nuclear genome
Bar labels: shared variants / all variants (percentage)
SNVs
2,960/2,976 (99.5%)
9,245/169,757 (5.4%)
Small indels
340/342 (99.4%)
1,034/21,543 (4.8%)
SVs
32/36 (88.9%)
191/4,649 (4.1%)
Variants shared by all five accessions (%)
