## Supplemental Methods for "Chromosome-scale comparative genomics reveals a genomic mosaic across the documented Hattan sake-brewing rice lineage"

**Supplementary Methods**

***Assembly cleanup and reference-guided scaffolding***

Before scaffolding, cleanup_short_contigs screened primary contigs shorter than 1 Mb for organelle-derived and rDNA-dominated sequences. Longer contigs passed unchanged through this step. Plastid- and mitochondrial-like contigs were identified against NC_001320.1 and NC_011033.1, respectively, and rDNA was identified using Barrnap v0.9. The workflow's default filtering thresholds were used. The retained contigs were scaffolded against AGIS1.0 using RagTag v2.1.0.

***Gene-model construction and representative-proteome preparation***

AGIS1.0/RAP-DB-supported protein-coding gene models were transferred to each cleaned, AGIS1.0-guided scaffold assembly using LiftOn v1.0.9. Partial mappings were excluded, copy recovery and coding-sequence-aware polishing were enabled, and a minimum sequence identity of 0.99 was applied. De novo gene predictions were generated for AGIS1.0 and the five Hattan assemblies using Helixer v0.3.6 with the land_plant lineage model, a window size of 100, an edge threshold of 0.1, a peak threshold of 0.99, and a minimum coding-length parameter of 60.

Transferred annotations were complemented with non-overlapping Helixer predictions using agat_sp_complement_annotations.pl from AGAT v1.7.0 without an additional AGAT minimum-size option. Following identifier normalization, a source-aware filter removed Helixer-derived transcripts encoding proteins shorter than 50 amino acids while retaining transferred and reference-derived models. CDS and protein sequences were extracted using gffread v0.12.8. The longest protein isoform associated with each gene was selected as its representative sequence and used for OrthoFinder v3.1.5. Subsequent header normalization did not alter amino-acid sequences.

BUSCO v6.0.0 was run in proteins mode on one representative protein per gene using poales_odb12, which comprised 6,282 orthologs. OMArk v0.4.1 was run on all annotated protein isoforms using OMAmer v2.1.2 and the LUCA database built from OMA All.Jul2024. Isoform-to-gene mappings were supplied so that OMArk selected the best-mapped isoform per gene for gene-level assessment. OMArk selected *Oryza* as the assessment lineage, comprising 15,087 conserved hierarchical orthologous groups. Summary metrics are provided in Supplementary Table S2.

***Genome-wide and local phylogenetic analyses***

The primary nucleotide alignment comprised 174,754 complete, homozygous, biallelic SNVs extracted from the repeat-filtered six-genome Cactus VCF. The independently generated DeepVariant-plus-Longshot alignment comprised 168,542 sites and was used for cross-pipeline validation. OrthoFinder v3.1.5 provided a concatenated representative-protein alignment of 11,056,173 residues for sequence-type validation.

Maximum-likelihood phylogenies were inferred using IQ-TREE v3.1.3. ModelFinder with ascertainment-bias correction was applied to the SNV-only alignments. The primary Cactus analysis selected TVM+F+ASC. Branch support was evaluated using 1,000 SH-aLRT replicates and 1,000 ultrafast-bootstrap replicates. Pairwise maximum-likelihood distances were obtained under the same fitted model. Trees were interpreted as unrooted. AGIS1.0 was placed at the base only for visualization and was not treated as a biological outgroup.

Chromosome-specific trees were evaluated for all 12 chromosomes using both ModelFinder and a fixed TVM+F+ASC sensitivity model. Chromosomes 8 and 12 were partitioned into non-overlapping 1-Mb windows, with terminal partial windows retained. A parsimony-informative site was defined as a biallelic SNV at which each allele was represented in at least two of the six genomes. Maximum-likelihood trees were inferred only for windows containing at least 50 parsimony-informative sites, yielding 17 windows on chromosome 8 and eight on chromosome 12. Cactus and DVLS classifications were compared in the focal regions, and cross-pipeline discordant segments were examined at 100-kb resolution.

***Genome-wide inheritance-mosaic classification and refinement of candidate transition intervals***

Reference positions aligned across all six genomes were extracted from the Cactus HAL alignment using HAL utilities from a validated Cactus v3.1.4 container. Bases overlapping the merged repeat mask were excluded. The resulting callable mask comprised 244,499,272 bp across the 12 AGIS1.0 nuclear chromosomes. Complete-case Cactus SNVs retained by this mask were summarized in non-overlapping 100-kb, 500-kb, and 1-Mb windows.

The 500-kb windows formed the inheritance-mosaic backbone. The 100-kb windows were used to refine category transitions, and the 1-Mb windows provided broad-scale support. Windows were classified from allele-direction-independent six-genome genotype patterns into prespecified lineage-event or accession-specific categories. The five Hattan positions in each genotype code were ordered as Hattan No. 10, Hattan No. 35, Hattan-nishiki No. 1, Hattan-nishiki No. 2, and Hattanso. At each biallelic SNV, 0 denotes the AGIS1.0 reference allele and 1 denotes the alternate allele. Complementary codes were treated as the same allele-direction-independent split. The callable fraction was calculated as the number of callable AGIS1.0 bases divided by the actual physical length of each window. Informative SNVs were complete-case homozygous biallelic Cactus SNVs within the callable mask and were not restricted to parsimony-informative sites. The dominant-pattern fraction was calculated as the number of SNVs in the most frequent allele-sharing category divided by the total number of these SNVs in the window, including those assigned to the other-pattern class. Assignment required a callable fraction of at least 0.20, at least three informative SNVs, a dominant-pattern fraction of at least 0.60, and a dominant pattern matching a prespecified category. Assignments with a callable fraction of at least 0.40 and at least five informative SNVs were designated high confidence. Assignments meeting only the minimum criteria were designated provisional. All remaining windows were unresolved. Consecutive windows with the same category were merged into blocks without bridging unresolved gaps.

Candidate transitions were recorded where the assigned category changed between adjacent blocks. The 100-kb classifications were used to delimit each transition by the nearest windows supporting different categories. These intervals represent changes in classified genomic state and were not interpreted as demonstrated molecular recombination breakpoints. Unresolved windows were further classified as mixed or complex, insufficiently callable, or insufficiently informative for supplementary reporting. Validation against DVLS required exact matches of chromosome, position, reference allele, and alternate allele. Missing DVLS genotypes were not imputed as reference. Category definitions and integrated block assignments, including unresolved subcategories, are reported in Supplementary Table S4.

***Comparison of Shuho SNV states with Hattan No. 10 and Hattan No. 35 using public sequence data***

Public whole-genome sequence data for the rice cultivar recorded as Shuho (listed as SHUHOU in the sequence archive) were obtained from DDBJ/NCBI run DRR054268. Associated records were experiment DRX049118, DRA sample DRS030033, BioSample SAMD00046017, DRA study DRP003133, and BioProject PRJDB4518. The Illumina HiSeq 2000 run had a paired-end layout and was associated with the japonica-rice sequencing project of Yano et al. (2016). The SRA Normalized Format archive was retrieved using SRA Toolkit v3.2.1 and converted using fasterq-dump --split-files without read trimming or length filtering. The archive yielded 3,868,427 read pairs, corresponding to 7,736,854 reads of 100 bp.

Reads were mapped to the six-genome pangenome graph using vg giraffe v1.66.0-44-gbfd254dcf with the corresponding GBZ, distance, minimizer, and zipcode indexes. BAM output was restricted to the 12 AGIS1.0 nuclear reference paths. Alignments were name-sorted, mate information was repaired, and the reads were coordinate-sorted and duplicate-marked using SAMtools v1.22. AGIS path prefixes in the BAM header were normalized to the chromosome accessions used for marker calling. The graph alignment contained all 7,736,854 reads. Of these, 7,334,228 reads, or 94.80%, mapped and 7,184,396 reads, or 92.86%, were properly paired.

The validation set comprised 25,833 biallelic SNVs at which the assembly-derived Hattan No. 10 and Hattan No. 35 genotypes carried different alleles. The Hattan No. 35 allele was defined as the expected state and the Hattan No. 10 allele as the contrast state. Allele-constrained calls were generated using BCFtools v1.22. mpileup required a minimum mapping quality of 30 and a minimum base quality of 20, skipped indels, and retained FORMAT/AD and FORMAT/DP. Multiallelic calling was constrained to the two specified marker alleles. Duplicate-marked reads were excluded under the pileup filtering defaults.

A marker was considered covered when an allele-depth field was available. Informative reads were those assigned specifically to the Hattan No. 35 or Hattan No. 10 allele. Reads supporting neither allele were excluded from the informative-read denominator. For each marker, the log10 likelihood ratio favoring Hattan No. 35 was calculated as (PL_H10/H10 − PL_H35/H35)/10. Positive values favored the Hattan No. 35 state. Marker and read summaries were interpreted descriptively because observations within linked genomic regions were not statistically independent.

Reference-state sensitivity was evaluated using markers at which the Hattan No. 35 allele was nonreference relative to AGIS1.0. Depth sensitivity was evaluated at minimum informative depths of one to five reads. Mapping-representation sensitivity was assessed by mapping the same read pairs to the 12 AGIS1.0 nuclear chromosomes using BWA-MEM2 v2.2.1. The alignments received the same SAMtools processing, and the unchanged marker set was called using the same BCFtools quality thresholds. The linear alignment contained all 7,736,854 primary reads. Of these, 7,494,578 reads, or 96.87%, mapped and 7,417,302 reads, or 95.87%, were properly paired.

Hattan No. 35-nonreference markers were assigned to non-overlapping 100-kb and 500-kb windows, and their marker-level log10 likelihood ratios were summed. Positive sums favored Hattan No. 35 and negative sums favored Hattan No. 10. Reported regional results were restricted to 500-kb windows containing at least five covered Hattan No. 35-nonreference markers. Mapping, marker-support, depth, and window-sensitivity summaries are provided in Supplementary Fig. S5 and Supplementary Table S5.

***Functional annotation of SNVs and small indels***

Complete, non-conflicting, biallelic SNVs and simple 1–50-bp insertions or deletions were extracted from the Cactus VCF. Indels of exactly 50 bp were assigned to this small-variant class and were excluded from final SV reporting. Variant-type-specific callability required the affected base for SNVs, the anchor and following base for insertions, and the external flanks for deletions. Variants were normalized against AGIS1.0 and annotated using snpEff v5.4c with a custom database constructed from the prepared AGIS1.0 genome and gene annotation. The database contained 42,013 genes and 46,644 protein-coding transcripts. All coding sequences passed the CDS check. A total of 105 transcripts with previously documented protein-sequence discrepancies were retained as flagged models.

Primary analyses were restricted to variants in lineage-assigned regions whose genotype partition agreed with the regional category. The most severe predicted effect was retained for each variant. Candidate HIGH-impact variants were evaluated for reference identity, callable status, gene-model quality, assembly context, and exact-allele DVLS support. When another primary-callable variant occurred within 10 bp, accession-specific homozygous compound haplotypes were reconstructed and reannotated. A candidate entered the final primary HIGH-impact set only when its HIGH consequence persisted after compound annotation, where applicable, and no unresolved technical or gene-model concern remained. The final classification indicates technical confidence and not phenotypic causality. Curated variants and their supporting evidence are reported in Supplementary Table S6.

***Structural-variant evidence integration***

Structural variants >50 bp were standardized in AGIS1.0 coordinates using three evidence families. Long-read evidence comprised consensus calls from Sniffles2 v2.7.0, cuteSV v2.1.2, and pbsv v2.11.0. Assembly evidence comprised whole-genome alignments generated using nucmer from MUMmer4 v4.0.1 and interpreted using SyRI v1.7.1. Graph evidence comprised explicit insertion and deletion alleles from the Cactus graph. Read-based SVIM, SVIM-asm, and legacy pseudochromosome SyRI results were used for sensitivity or caller-concordance checks and were not counted as independent evidence families.

Within each Hattan accession, insertions were matched by reciprocal-best assignment when breakpoint distance was no greater than 500 bp and the size ratio was at least 0.70. Deletions and inversions additionally required both breakpoints to be within 500 bp, reciprocal overlap of at least 0.60, a size ratio of at least 0.70, and reciprocal-best matching. Read BND consensus required matching chromosome pairs, both breakpoints within 1 kb, reciprocal-best assignment, and support from at least two callers. SyRI translocations were linked to read BNDs when both breakpoints were within 5 kb and projection width was no greater than 20 kb. Across Hattan accessions, reciprocal-best matches were clustered using complete linkage, such that every pair of events within a cluster satisfied the matching thresholds. Each cluster was restricted to at most one event per accession, preventing transitive chaining of nearby but distinct events. Integrated events were annotated with gene, repeat, projected query-gap, and inheritance-mosaic context.

An across-accession event entered the primary multi-evidence set when at least one accession-specific member was supported by two or more independent evidence families. Events overlapping or located within 1 kb of a high-confidence projected query-assembly gap in a Hattan accession in which the event was detected were deferred for review. Events located 1–10 kb from such gaps were retained with a caution flag. Replicated events lacking a second independent evidence family were retained separately as single-family events. PAV information was treated as auxiliary and was not counted as an independent evidence family or used to increase SV confidence. Detection patterns summarized observed calls across Hattan accessions and were not interpreted by themselves as proof of biological absence. Curated events and their evidence are reported in Supplementary Table S7.

***Pangene construction and gene presence–absence variation curation***

A coordinate-first pangene matrix was generated using a modified get_pangenes workflow, version 27052025, with a minimum GFF length of 60, P=95, and N=0. Whole-genome alignments and gene coordinates were used separately for the 12 nuclear chromosomes. Initial assignments were reconciled with OrthoFinder v3.1.5 representative-protein orthogroups. Screening identified 1,732 rows matching breeding-step or accession-specific presence–absence patterns among 35,639 initial pangene rows. Candidates were prioritized when a LiftOn-derived source model had compatible protein and gene-model evidence without critical technical or tandem-context flags. Prioritization additionally required direct-SV support with an exact, compatible, or partial accession pattern, or positional-copy evidence. These prioritized records underwent the locus-level exclusion and reconciliation described below. Orthogroup membership was used to guide correspondence rather than as a conclusive test of local gene absence.

Candidates identified in a Hattan assembly were projected to AGIS1.0 coordinates using the Cactus HAL alignment. For prioritization, insertion and deletion evidence was searched within 5 kb of the projected reference interval, and event detection patterns were compared with the candidate pattern. Assembly gaps within 1 kb, chromosome ends within 100 kb, pseudogene models, and same-orthogroup copies within 100 kb were flagged for technical or tandem-context review. Protein homology was evaluated using Miniprot v0.18-r281 across all six genomes. Initial intact-like hits required query coverage ≥0.80, identity ≥0.70, and no reported frameshift or internal stop. Partial or disrupted-like hits required coverage ≥0.50 and identity ≥0.50. These searches identified corresponding sequences and related copies for subsequent locus evaluation. A nearby SV was not by itself treated as proof of whole-gene gain or loss.

Locus correspondence was evaluated independently of orthogroup membership using surrounding shared genes, DNA alignments generated with Minimap2 v2.31-r1302, and local protein alignments. Up to four shared flanking genes and contexts extending up to 500 kb on either side were used to distinguish the focal locus from related copies. At a resolved corresponding locus, ≥95% source-CDS alignment with nucleotide identity ≥85% supported retained sequence, while >10% but <95% CDS alignment at identity ≥85% was recorded as partial sequence. Alignment of ≤10% of the source CDS identified a local-absence candidate requiring structural and read confirmation. Short query proteins that aligned within a longer retained model were not interpreted as whole-gene absence. Source models were checked by direct CDS translation and comparison with the representative-protein FASTA. These alignment thresholds guided sequence review; they were not treated as proof of equivalent gene structure or function. Confidently retained full-length counterparts and coding/gene-model differences were excluded from the PAV candidate list, whereas partial-sequence candidates and cases requiring further evaluation were retained.

Structural evidence was checked using the material-specific alignment intervals, including the query-side coordinates of SyRI events. A gene-deleting event had to cover at least 90% of the source CDS in reference coordinates and lack a substantially matching protein counterpart (query coverage ≥0.80 and identity ≥0.70) in the query replacement interval. The effect on the gene in each accession was evaluated separately from the detection pattern of the across-accession SV group. Compatible genes sharing the same actual gene-deleting event and presence–absence pattern were subsequently grouped as one independent PAV locus.

HiFi reads overlapping the candidate regions in either existing AGIS1.0 alignment were combined and deduplicated by read name. These reads were remapped to each complete corresponding nuclear assembly, including unplaced sequences, using Minimap2 v2.31-r1302 with the map-hifi preset, --eqx, --secondary=yes, and -N20. SAMtools v1.22 was used for alignment processing. Read evaluation required mapping quality ≥30, with base quality ≥20 for sequence-depth summaries. Support for the assembled local allele required at least 90% breadth at depth ≥3 and either at least three templates spanning the allele with ≥300 matched bases on each side, or multi-family structural support together with at least three qualifying templates at each boundary. Read coverage alone was insufficient to resolve repeated or organelle-like sequences.

Highly similar copies elsewhere were searched without requiring the same orthogroup. Copy-retention evidence required protein-query coverage ≥95%, identity ≥95%, and no reported frameshift or internal stop. Local absence with such a copy was recorded as positional PAV. Among the remaining supported loci, those with gene-deleting events supported by at least two evidence families were recorded as direct structural PAVs; other supported loci were recorded as locally read-validated PAVs. These mutually exclusive reporting classes distinguish copy context and the supporting evidence. Positional PAVs can themselves have multi-family structural support.

Correspondence between alternative candidate records was accepted only with reciprocal sequence and positional support. Corresponding records were merged before counting candidates; candidates with partial correspondence between alternative source models were retained as one unit for further evaluation. Candidates were also retained when locus or copy correspondence, assembly or gene-model quality, or direct read support required further evaluation. For LOC107276587, Hattanso showed 97.51% source-CDS alignment at 85.25% identity without a full-length protein counterpart in the existing search. This borderline correspondence was retained for further evaluation and recorded as unresolved in the candidate table. Neither a missing model nor an unresolved correspondence was converted into sequence absence. Supported patterns outside the prespecified breeding-step or accession-specific categories were reported separately from the primary lineage-associated PAV set.

Supplementary Table S9 lists one record per reconciled PAV candidate unit. It includes the 46 supported PAVs and candidates with partial sequence or unresolved evidence, with supporting observations in the same table. Retained, Absent, Partial, and Unresolved describe local DNA correspondence; an Absent state alone does not indicate that all read-support criteria were met. All 85 PAV candidate units were plotted at their projected AGIS1.0 positions in Figure 5C, distinguishing the 46 supported PAVs from the 39 candidates requiring further evaluation. The supported primary lineage-associated set comprises 44 gene units across 36 independent loci and was used for the supported-locus chromosome summaries, TE analyses, and PAV enrichment targets. Genes sharing one structural event and presence–absence pattern share a primary PAV locus ID. Earlier candidate records and their correspondence to the reconciled units were retained in the analysis records for reproducibility.

***Transposable-element annotation and spatial relationships with SVs, PAV loci, and candidate transition intervals***

HiTE v3.3.2 annotations were available for AGIS1.0 and all five Hattan assemblies. EDTA v2.2.2 annotations were available for the five Hattan assemblies. AGIS1.0 HiTE intervals provided the primary reference-coordinate TE track. Query-side HiTE and EDTA annotations were used for locus-level validation. RepeatModeler v2.0.6 and RepeatMasker v4.1.9 were used as auxiliary sensitivity resources.

Primary tests evaluated breakpoints from high-confidence SVs supported by at least two evidence families. Secondary target sets comprised gene-disrupting SVs, primary lineage-associated PAV boundaries, and candidate transition intervals. For PAV loci, the median reference endpoints of confirmed gene-deleting SVs were used when available; otherwise the evaluated gene-interval endpoints were used and recorded as such. Exploratory tandem-context targets represented candidate-region context rather than confirmed PAVs. Additional subsets were treated as exploratory and are identified in Supplementary Table S8. TE overlap and proximity within 100 bp, 1 kb, and 10 kb were calculated for each target set. Overlap required strict intersection of half-open target and TE intervals. Intervals that met only at an endpoint were not scored as overlapping. Proximity was the minimum gap to the primary TE track. For each source event or locus, the minimum distance across its component intervals was used, and an entity was scored as overlapping when any component interval overlapped a TE.

Null distributions were generated by holding the AGIS1.0 TE track fixed and applying one common random offset to all targets on each chromosome. Offsets were sampled independently among chromosomes, and intervals that crossed a chromosome end were wrapped to the other end. This retained chromosome assignment, interval length, paired boundaries, and within-chromosome clustering. Primary union tests used 10,000 permutations, and class-specific tests used 2,000 permutations. SV-stratified and TE-definition sensitivity analyses used 3,000 permutations. Global and local null-model comparisons used 5,000 permutations. The local model sampled signed shifts with absolute values from 100 kb to 5 Mb to assess sensitivity to broad variation in TE density.

Empirical one-sided probabilities used a plus-one correction. Two-sided probabilities were twice the smaller one-sided probability, capped at one. P values were adjusted using the Benjamini–Hochberg procedure within prespecified test families, including union analysis unit, class-specific metric, and sensitivity-analysis dimension or null model and metric. Significant proximity was interpreted as spatial association and not as evidence that a TE caused the structural change. PAV targets were updated from the supported locus set, and the shared BH test families were recalculated. Existing SV-stratified, null-model, and TE-definition sensitivity estimates were retained. Primary SV results are shown in Supplementary Fig. S6. Complete results for all target sets and sensitivity analyses are provided in Supplementary Table S8.

***Functional annotation of six rice proteomes and enrichment analysis of lineage- and variant-associated gene sets***

Functional annotations generated using EnTAP v2.3.0 were reconciled across the six proteomes. GO identifiers were normalized to the 2026-06-19 archive with internal data version 2026-06-15. is_a and part_of ancestors were propagated. Direct annotations and annotations supported in at least two genomes were retained as sensitivity models. Fixed EnTAP and eggNOG-mapper v2.1.12 assignments supplied KEGG identifiers. KEGG release 119.0 was used to validate identifiers, obtain current labels, and define a plant-focused *Oryza sativa* japonica pathway scope.

One curated pangene unit was treated as one statistical unit. Six pairs of rows confirmed to represent corresponding gene loci were merged, yielding 35,633 units while preserving the original gene-model IDs. The primary functional background excluded 32 rows associated with unresolved correspondence, leaving 35,601 units; a sensitivity analysis retained these rows. Prespecified targets comprised genes in lineage-assigned intervals, final primary HIGH-impact variant genes, multi-evidence SV-overlapped genes, and supported PAV genes. PAV enrichment targets were restricted to the primary lineage-associated set and comprised all 44 gene units in this set, the 40 with multi-family gene-deleting SV support, and the 18 with such support and no detected close full-length copy elsewhere. Each test used the functionally annotated units in the corresponding background.

At least ten annotated target units were required for testing. Sets below this threshold were recorded as not tested rather than as nonsignificant. One-sided hypergeometric tests were followed by Benjamini–Hochberg correction within each target set and ontology or entity class. Reported GO terms required FDR ≤0.05 and at least three target units. Reported pathways additionally required membership in the current rice pathway list and the predefined plant-focused hierarchy. Significant GO–pathway pairs were tested for shared pangene units within target sets. Reportable pairs required at least two shared units and FDR ≤0.05. Cross-ontology overlap was interpreted as concordance and not as independent replication. Complete target-set and sensitivity summaries are provided in Supplementary Table S11.

***Candidate-locus review***

Fifteen candidate loci were reviewed using structural evidence, inheritance-pattern agreement, gene overlap and model quality, transposable-element context, and functional evidence. This review was intended to summarize locus-level support and limitations rather than test phenotype associations. Evidence for all 15 loci is reported in Supplementary Table S10, with links to the retained PAV candidate units in Supplementary Table S9 and an indication of local PAV support. The presence of a related SV was distinguished from confirmation of whole-gene PAV.

**Nonreference SNVs shared across the Hattan lineage**

Shared SNVs were selected from the existing complete-case, homozygous, biallelic Cactus dataset by requiring the same alternate allele in the sampled Hattanso and all four derived cultivars, with AGIS1.0 carrying the reference allele. Only sites retained by the six-genome aligned, nonrepeat callable mask were included. The callable mask spans 244,499,272 bp across the 12 AGIS1.0 nuclear chromosomes and contains 172,733 complete-case SNVs. The 174,754-site phylogenetic alignment has a different callable-region scope. Shared-SNV extraction yielded 12,205 sites.

Chromosome densities were calculated as shared-SNV counts divided by callable bases and multiplied by 1,000,000. Density ratios used the genome-wide mean of 12,205/244,499,272 × 1,000,000. Non-overlapping 100-kb and 500-kb grids were reused from the mosaic analysis. Window-density rankings were restricted to 500-kb windows with a callable fraction of at least 0.40. The existing lineage-shared mosaic blocks supplied the region assignments. The 100-kb summaries describe local variation within these blocks. Site counts and density ratios were treated descriptively because nearby variants are linked.

The independently generated DVLS strict common VCF for each accession was queried at the shared sites. Support required an exact chromosome, position, reference-allele, and alternate-allele match, FILTER=PASS, and a homozygous-alternate genotype in each of the five accessions. Missing records were retained as missing evidence. The number of supporting accessions was recorded per site, and the all-five-supported subset was summarized by chromosome and window using the same callable denominators. Of the 12,205 shared SNVs, 7,686 met the all-five DVLS support criterion, including 1,143 in Chr4:17.5–18.5 Mb. The 18.0–18.5 Mb window retained the highest shared-SNV density when only the all-five DVLS-supported subset was considered. Supplementary Table S12 contains chromosome, window, and site-level support summaries.

**Shared small indels and structural variants in the Chr4 region**

The existing six-genome complete-case Cactus VCF within the primary callable mask also supplied small insertions and deletions. Sharing required the same nonreference allele in all five Hattan accessions. The complete small-variant set comprised 172,733 SNVs and 21,885 small indels, including 12,205 shared SNVs and 1,374 shared small indels. Variant positions were summarized in AGIS1.0 coordinates. The focal interval was the existing lineage-shared mosaic block at Chr4:17,500,000–18,500,000 on the 0-based, half-open coordinate system.

Structural comparisons used the existing set of 4,685 events classified as TIER1_HIGH_CONFIDENCE_MULTI_FAMILY after structural-annotation filtering. A shared SV had the across-accession detection pattern 11111. This definition identifies matched event detections; full inserted-sequence identity was not required. Multi-family support in every accession was recorded separately. Of 223 shared events genome-wide, 151 met this stricter per-accession requirement. SVs were assigned to windows by their left AGIS1.0 reference anchor; alternative end, midpoint, either-boundary, and interval-overlap assignments were also examined. SV denominators comprise the observed multi-evidence events, whereas the small-variant denominators comprise complete-case callable sites.

Local profiles used non-overlapping 100-kb windows across Chr4:16.5–19.5 Mb. Shared/all fractions were calculated separately for each variant class inside Chr4:17.5–18.5 Mb and in the remaining nuclear genome. For the focal interval, the primary and strict single-copy callable masks both contained 525,547 bp. The focal counts were unchanged by the alternative SV coordinate rules. A comparison restricted to long-read plus assembly-WGA SV evidence retained 14 shared events among 18 focal events. Window counts, focal-event evidence, coordinate sensitivity, and the shared small-variant and SV catalogues are provided in Supplementary Table S12. These comparisons summarize linked genomic variation descriptively.

*OsWAK43* and *OsWAK44* were identified by the RAP-DB identifiers Os04g0368000 and Os04g0368300 and the corresponding AGIS1.0 models LOC9269553 and LOC107275485. Their positions are shown in Supplementary Fig. S8B. Published candidate-gene and marker information was checked against Lafarge et al. (2017). Local flanking sequences placed the *OsWAK43*-associated marker D04_17876533 at AGIS1.0 Chr4:18,225,987. AGIS1.0 and all five Hattan assemblies carried A at this marker. Gene-coordinate mappings and marker states are reported in Supplementary Table S12.

**Sequence and read validation of the shared *SSIIb* allele**

*SSIIb* was identified as Os02g0744700 and AGIS1.0 gene LOC4330709. The final gene model for each assembly was used to extract the coding sequence, which was translated and compared with the corresponding protein sequence used in the original annotation. The AGIS1.0 G>A variant at NC_089036.1:31,729,214 maps to c.1756C>T on the reverse-strand coding sequence and p.Leu586Phe. Local 40-bp flanks on each side anchored the site in each assembly and at IRGSP-1.0 Chr2:31,234,046.

Existing HiFi BAM alignments produced with minimap2 and pbmm2 were evaluated separately. Targeted alignments were extracted with SAMtools v1.22, and bases were read from the SAM CIGAR and base-quality fields without BAQ adjustment. The primary analysis required mapping quality ≥30 and base quality ≥20 and excluded unmapped, secondary, supplementary, QC-failed, and duplicate-marked alignments. Each read template was counted once, and templates with discordant qualifying bases were discarded. Reference, alternate, other-base, and discordant-template counts were recorded. Alternate-allele support was summarized as alternate templates divided by quality-filtered templates. A sensitivity analysis used mapping quality ≥20 and base quality ≥10. Both alignments use the same HiFi data. Exact-allele PASS homozygous-alternate records in the existing DVLS strict VCFs provided an additional callset comparison.

The corresponding IRGSP-1.0 site was examined in the 1,358 public material records available through RAP-DB/TASUKE+ on 4 November 2026 (Kawahara et al. 2026). Raw depth and SNP calls filtered at QUAL ≥30 and DP ≥10 were retrieved separately. Explicit homozygous SNP genotypes passing these filters were distinguished from records with no variant entry but depth ≥10, which were classified as reference-compatible. Heterozygous, low-depth, and failed-filter records were excluded from the homozygous/reference-compatible denominator. TEJ denotes the database temperate-japonica group. Fractions describe database records, including repeated cultivar names. Public accessions, depths, and call origins are reported in Supplementary Table S13.
